# Model-based prediction of spatial gene expression via generative linear mapping

**DOI:** 10.1101/2020.05.21.107847

**Authors:** Yasushi Okochi, Shunta Sakaguchi, Ken Nakae, Takefumi Kondo, Honda Naoki

## Abstract

Decoding spatial transcriptomes from single-cell RNA sequencing (scRNA-seq) data has become a fundamental technique for understanding multicellular systems; however, existing computational methods lack both accuracy and biological interpretability due to their model-free frameworks. Here, we introduced Perler, a model-based method to integrate scRNA-seq data with reference *in situ* hybridization (ISH) data. To calibrate differences between these datasets, we developed a biologically interpretable model that uses generative linear mapping based on a Gaussian-mixture model using the Expectation-Maximization algorithm. Perler accurately predicted the spatial gene expression of *Drosophila* embryos, zebrafish embryos, mammalian liver, and mouse visual cortex from scRNA-seq data. Furthermore, the reconstructed transcriptomes did not over-fit the ISH data and preserved the timing information of the scRNA-seq data. These results demonstrated the generalizability of Perler for dataset integration, thereby providing a biologically interpretable framework for accurate reconstruction of spatial transcriptomes in any multicellular system.

## Introduction

Genes are heterogeneously expressed in multicellular systems, and their spatial profiles are tightly linked to biological functions. In developing embryos, spatial gene-expression patterns are responsible for coordinated cell behavior (e.g., differentiation and deformation) that regulates morphogenesis^1^. Additionally, within organ tissues, cells at different locations play different roles in organ function based on their gene-expression patterns^2^. Thus, identification of spatial genome-wide gene-expression profiles is key to understanding the functions of various multicellular systems. *In situ* hybridization (ISH) has been widely used to visualize spatial profiles of gene expression; however, application of this method is generally limited to only small numbers of genes. By contrast, the single-cell RNA sequencing (scRNA-seq) method developed during the previous decade has enabled measurement of genome-wide gene-expression profiles in tissues at the single-cell level^3^. However, this method requires tissue dissociation, which leads to loss of spatial information for the original cells.

To compensate for the lost spatial information, new computational approaches have emerged (Seurat v.1^4^, DistMap^5^, Achim et al.^6^, Halpern et al.^7^), enabling reconstruction of genome-wide spatial expression profiles from scRNA-seq data by integrating existing ISH data as a spatial reference map *in silico*. However, their methods require binarization of gene-expression data^8^, which leads to unsatisfactory accuracy, or tissue-specific modelling, which leads difficulty in application to other systems. Recently, the seminal methods Seurat (v.3)^9^ and Liger^10^ were developed to address gene-expression data as continuous variables in a non-tissue-specific manner. These methods match the distributions of ISH and scRNA-seq data points by using dimensionality reductions [e.g., canonical correlation analysis (CCA)]^11^ and integrative non-negative matrix factorization (iNMF)^12^, followed by mapping the scRNA-seq data points to the nearest ISH data points according to Euclidean distance using Nearest-Neighbor (NN) methods (e.g., k-NN^13^ and mutual NN^14^). However, a major issue is that the flexibility of the methods allow mapping of ISH data to scRNA-seq data without any models of the underlying scRNA-seq data structure. Specifically, these methods do not account for difference in gene-expression noise associated with each gene. Given this model-free property, these methods are dependent upon nonlinear NN mapping, which innately causes over-fitting to the reference ISH data.

To address these issues, we propose a novel model-based computational method for probabilistic embryo reconstruction by linear evaluation of scRNAseq (Perler), which reconstructs spatial gene-expression profiles via generative linear modeling in a biologically interpretable framework. Perler addresses gene-expression profiles as continuous variables and models generative linear mapping from ISH data points into the scRNA-seq space. To estimate parameters of the linear mapping, we developed a method based on the Expectation–Maximization (EM) algorithm^13^. Using the estimated parameters, we also propose an optimization method to infer spatial information of scRNA-seq data within a tissue sample. We applied this method to existing *Drosophila* scRNA-seq data^5^ and successfully reconstructed spatial gene-expression profiles in *Drosophila* early embryos that were more accurate than those generated using another spatial reconstruction method (DistMap^5^). Additionally, we showed that Perler can reconstruct a spatial gene-expression pattern that could not be fully predicted using previous methods, including Seurat (v.3), Liger, and DistMap. Further analysis revealed that Perler was able to preserve the timing information of the scRNA-seq data without over-fitting to the reference ISH data. Furthermore, we demonstrated that this method accurately predicted spatial gene-expression profiles in early zebrafish embryos^4^, the mammalian liver^7^, and the mouse visual cortex^15,16^. These findings demonstrate Perler as a robust, generalized framework for predicting spatial transcriptomes from any type of ISH data for any multicellular system without overfitting to the reference.

## Results

### Framework of spatial reconstruction in Perler

Perler is a novel computational method for model-based prediction of spatial genome-wide expression profiles from scRNA-seq data that works by referencing spatial gene-expression profiles measured by ISH (**Fig. 1a**). In general, scRNA-seq data have higher dimensionality (on the order of ∼10,000 genes) but does not contain information of spatial coordinates in tissues. By contrast, reference ISH data contain expression information for *D* genes in each cell or tissue subregion, with these referred to as landmark genes (e.g., *D* = 84 in *Drosophila melanogaster* early embryos) and tagged with spatial coordinates in tissues.

**Figure 1:**
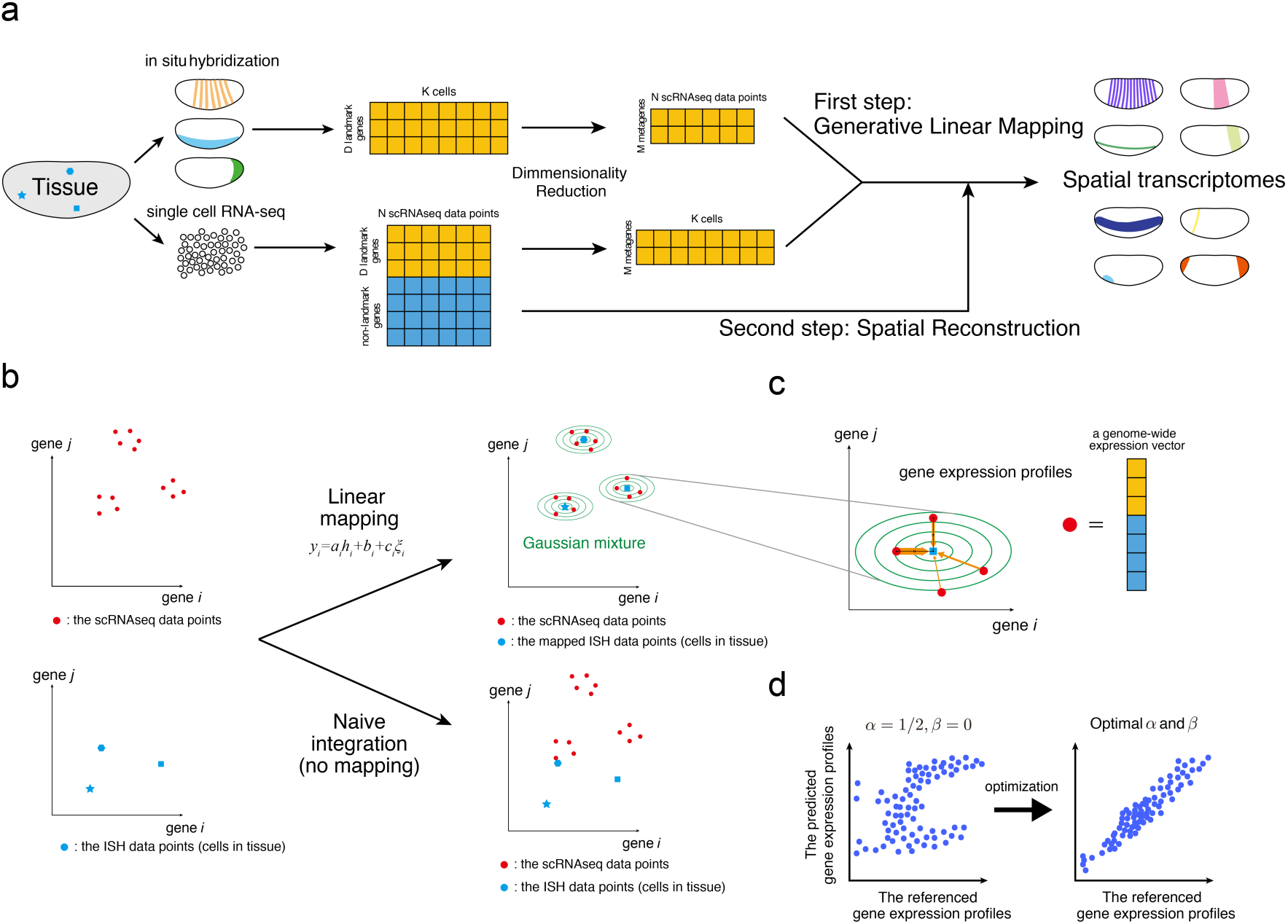
Schematic illustration of Perler. (a) Flow of data processing. (b) Generative linear mapping from ISH data to the scRNA-seq space. The left and right panels indicate scatter plots in high-dimensional ISH and scRNA-seq spaces. Because ISH data points did not match scRNAseq data points (Naïve integration) in the absence of mapping, ISH data points were mapped in order to fit the scRNA-seq data points the best using the EM algorithm. Blue points indicate ISH data points, red points indicate scRNA-seq data points, and green lines indicate contours of the estimated multivariate Gaussian distribution (see Methods). (c) Reconstruction/prediction of gene expression by Mahalanobis’ metric-based weighting (see Methods). Orange arrows indicate weights between scRNA-seq data points to cell *k*, and their widths reflect the Mahalanobis’ metric-based weights. Note that although two scRNAseq data points are equally distant from the ISH data point in the Euclid metric (black line), the weights are different. (d) Weight determination. The hyperparameters of the weighting function, *α* and *β*, are determined by cross-validation to ensure that the referenced gene-expression profiles correlate well with the predicted gene-expression profiles (see Methods). Dots correspond to cells in tissue. The left and right panels indicate the conceptual scatter plots of the expression levels of the genes before (*α* = 1/2, *β* = 0) and after parameter optimization, respectively.

The Perler procedure involves two steps. The first step estimates a generative linear model-based mapping function that transforms ISH data into the scRNA-seq space, thereby enabling calculation of pairwise distances between ISH data and scRNA-seq data (**Fig. 1b**). The second step reconstructs spatial gene-expression profiles according to the weighted mean of scRNA-seq data, which is optimized by the mapping function estimated in the first step (**Fig. 1c**).

The first step considers gene-specific differences between scRNA-seq and ISH measurements. For example, we assume that some genes are more or less sensitive to ISH or scRNA-seq and subject to high or low background signals in the associated data. We account for gene-specific noise intensity, because gene expression fluctuates over time in a gene-specific manner^17^. These differences in sensitivity, background signals, and noise intensity can be expressed by linear mapping:

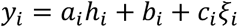

where *y*_*i*_ and *h*_*i*_ denote the expression levels of landmark gene *i* measured by scRNA-seq and ISH, respectively; *a*_*i*_, *b*_*i*_, and *c*_*i*_ are constant parameters of gene *i* and interpreted as the sensitivity coefficient, background signal, and noise intensity, respectively; and *ξ*_*i*_ indicates standard Gaussian noise. Note that *a*_*i*_, *b*_*i*_, and *c*_*i*_ are different for each gene, and that these parameter values are unknown. To estimate this linear mapping from the data, we developed a generative model in which scRNAseq data points are generated/derived from each cell in the tissue, whose expression is measured by ISH (see Methods). We then derived a parameter-estimation procedure based on the EM algorithsm (see Methods). Using the estimated parameters, a gene-expression vector for each cell in a given tissue sample measured by ISH can be mapped to the scRNA-seq space, thereby allowing evaluation of pairwise distances between ISH and scRNA-seq data.

The second step reconstructs the spatial gene-expression profile in tissue from scRNA-seq data. We estimated gene expression of each cell in a tissue sample according to the weighted mean of all scRNA-seq data points, where the weights were determined by the pairwise distances between cells in tissue samples measured using ISH and scRNA-seq data points (**Fig. 1c**). For the best prediction, we optimized the hyperparameters of the weighting function to ensure that the predicted and referenced landmark gene-expression profiles were well-correlated by cross-validation (**Fig. 1d**; see Methods). We then predicted non-landmark gene expression using the optimized weighting function.

Data were preprocessed before the first step. Some landmark genes redundantly exhibit similar spatial expression patterns, which can lead to biased parameter estimation and cause a loss of mapping ability. To reduce redundancy in scRNA-seq and ISH data, we performed dimensionality reduction using partial least squares correlation analysis (PLSC)^18^ (see Methods). Each factor in the reduced dimension can be interpreted as a “metagene”, which is representative among a highly correlated gene cluster, with its coordinate corresponding to the expression level of the metagene. In Perler, we regarded the metagene-expression level *i* (i.e., factor *i*) in the scRNA-seq and ISH spaces as *y*_*i*_ and *h*_*i*_ in the equation above (see Methods).

### Model-based mapping between scRNA-seq and ISH data

Previously, Karaiskos *et al*.^5^ measured gene expression in individual cells dissociated from early *D. melanogaster* embryos at developmental stage 6 by scRNA-seq, followed by development of a computational method (DistMap) to reconstruct the spatial gene-expression profile of the embryos from the scRNA-seq data. They used as reference data a spatial gene-expression atlas provided by the Berkeley *Drosophila* Transcription Network Project (BDTNP)^19, 20^, in which the expression of 84 landmark genes was quantitatively measured by fluorescent (FISH) at single-cell resolution at developmental stage 5.

In the present study, we applied Perler to the same scRNA-seq dataset and used the 84 landmark genes from the BDTNP atlas as the spatial reference map. We then predicted the spatial gene-expression profiles for 8840 non-landmark genes. To compare Perler results with those of DistMap, we used the same normalization methods for the scRNA-seq dataset as the previous study^5^. For preprocessing, we manually extracted 60 metagenes as non-redundant clusters of the landmark genes by dimensionality reduction and estimated the parameters of the linear mapping by integrating the scRNA-seq data with the ISH data (**Supplementary Fig. 1a–c**). The mapped ISH data points according to the linear mapping were distributed consistently with the scRNA-seq data points (**Fig. 2a, Supplementary Fig. 1d**). Additionally, the reconstructed and referenced gene-expression profiles were well-correlated following optimization of the hyperparameters (**Fig. 2b**). We then derived a posterior probability that a scRNA-seq data point was generated from each cell in the tissue sample and confirmed that the majority of scRNA-seq data points (82.7%) was assigned to three cells in the tissue with high confidence (>0.8) (**Fig. 2c, Supplementary Fig. 1e**). Moreover, we showed that the scRNA-seq data points were specifically assigned to cells in a small region (a few cell diameters) of the tissue (**Fig. 2d, Supplementary Fig. 1f and g**). The reconstruction accuracy of Perler (average Correlation Coefficient (aCC) = 0.76) was significantly higher than that of Seurat (v.3; aCC = 0.61), Liger (aCC = 0.61), and DistMap (aCC = 0.56) (**Fig. 2e and f, Supplementary Fig. 2**). These results demonstrated that Perler accurately reconstructed the spatial expression profiles of the landmark genes using ISH data of all the landmark genes for training and was capable of doing this via simple linear mapping.

**Figure 2:**
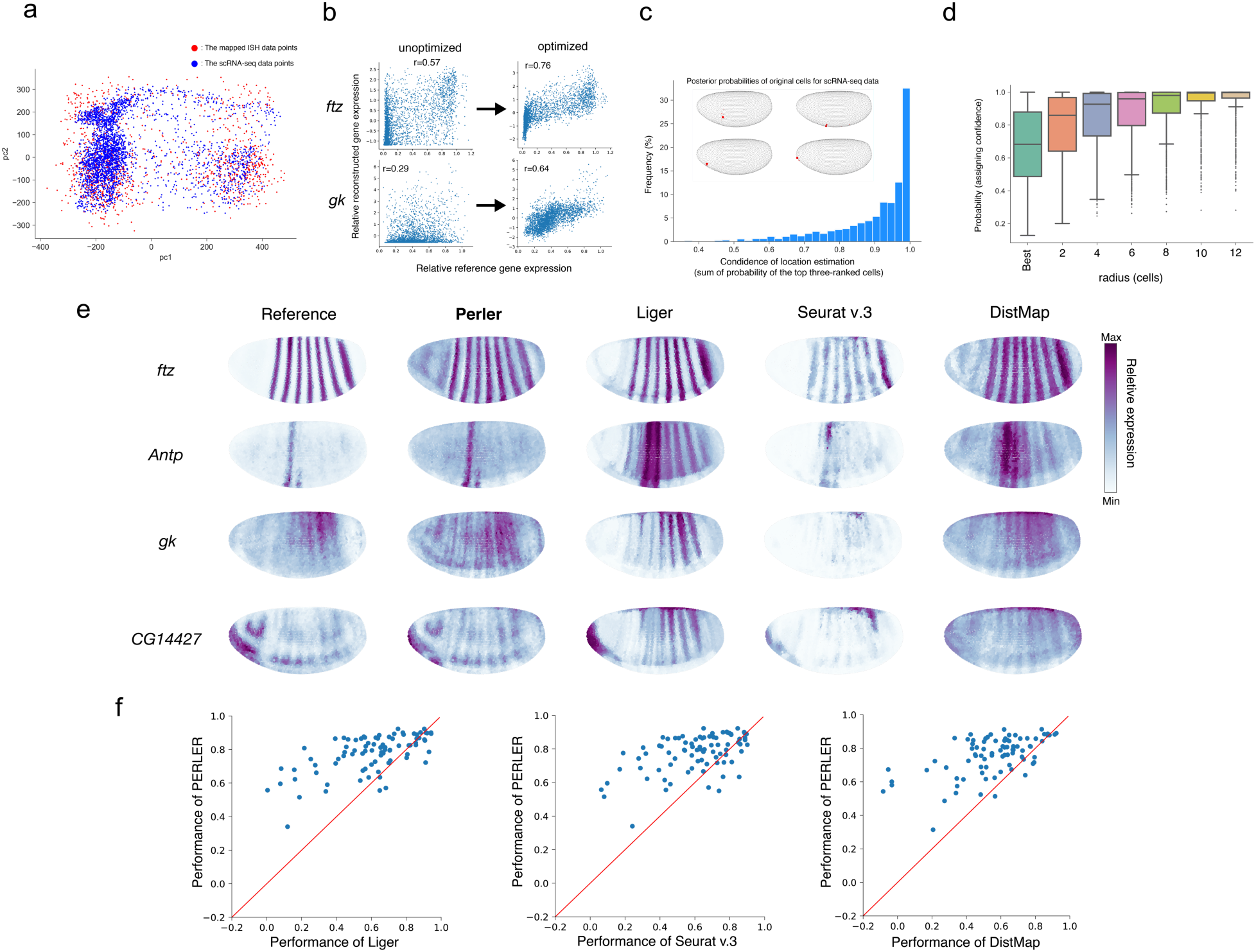
Generation of spatial gene expression profiles. (a) Scatter plot of mapped gene expression and scRNA-seq observations (Fig. 1b, upper right panel). Principal component analysis^13^ was used to visualize high-dimensional gene expression data into two dimensions. (b) Improved correlation between predicted and referenced data in the scRNA-seq space by optimizing the weighting function. (c) Histogram of the assigned confidence calculated as the posterior probabilities of the top three cells for each scRNA-seq data point. Perler assigned the majority of scRNA-seq data points (82.7%) to the top three cells with high confidence (>0.8). Examples of estimated original positions of each scRNA-seq data point (inset). Embryos are colored according to the posterior probabilities for the scRNA-seq data points. Point sizes indicate the magnitude of each posterior probability. Points with posterior probabilities below 0.001 were omitted. (d) Boxplot of the assigned specificity calculated as the posterior probabilities of circular regions for each scRNA-seq data point according to radius, with the center of each region representing the best assigned location for each data point. The boxplot has whiskers with a maximum 1.5 interquartile range, with black points indicating outliers. The radius was calculated by path length on the k-NN graph comprising all cells in the tissue (k = 6). (e) Reconstructions of the landmark genes by Perler, Liger, Seurat (v.3), and DistMap. (f) Comparison of Perler reconstruction performance with Liger (left, two-sided Wilcoxon test: p = 1.2×10^−9^), Seurat (v.3) (middle, two-sided Wilcoxon test: p = 1.9×10^−9^), and DistMap (right, two-sided Wilcoxon test: p = 1.4×10^−13^). Each dot indicates the reconstruction accuracies for each gene by Perler and other methods.

### Predictive ability of Perler

We then evaluated the predictive performance of Perler by conducting leave-one-gene-out cross-validation (LOOCV) in order to confirm whether gene expressions can be predicted following removal of the landmark gene of interest from the ISH data prior to training (**Supplementary Figs. 3 and 4**). The predictive accuracy of Perler (aCC = 0.59) was significantly higher than that of Seurat (v.3; aCC = 0.55), Liger (aCC = 0.53), and DistMap (aCC = 0.44) (**Supplementary Fig. 3**). Moreover, the predictive accuracy can be further improved by introducing maximum a posteriori (MAP) estimation at the M step in the EM algorithm (see Methods).

**Figure 3:**
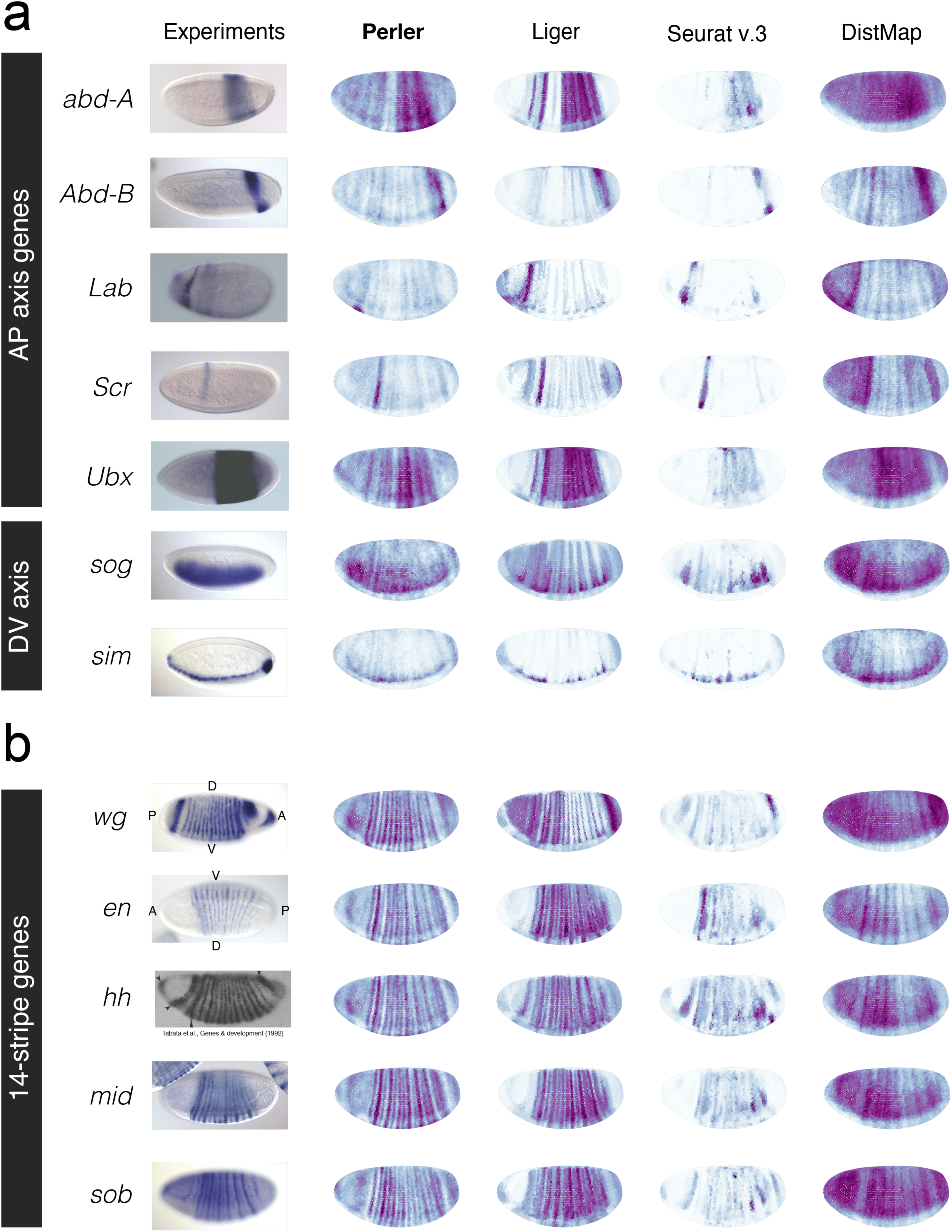
Spatial prediction of non-landmark genes. Predictions of non-landmark gene expression showing (a) spatial expression along the A–P and D–V axes and (b) a 14-stripe pattern according to Perler, Liger, Seurat (v.3), and DistMap. ISH image of *hh* reprinted from Tabata *et al*.^22^ under a Creative Commons License (Attribution: Non-Commercial 4.0 International License).

**Figure 4:**
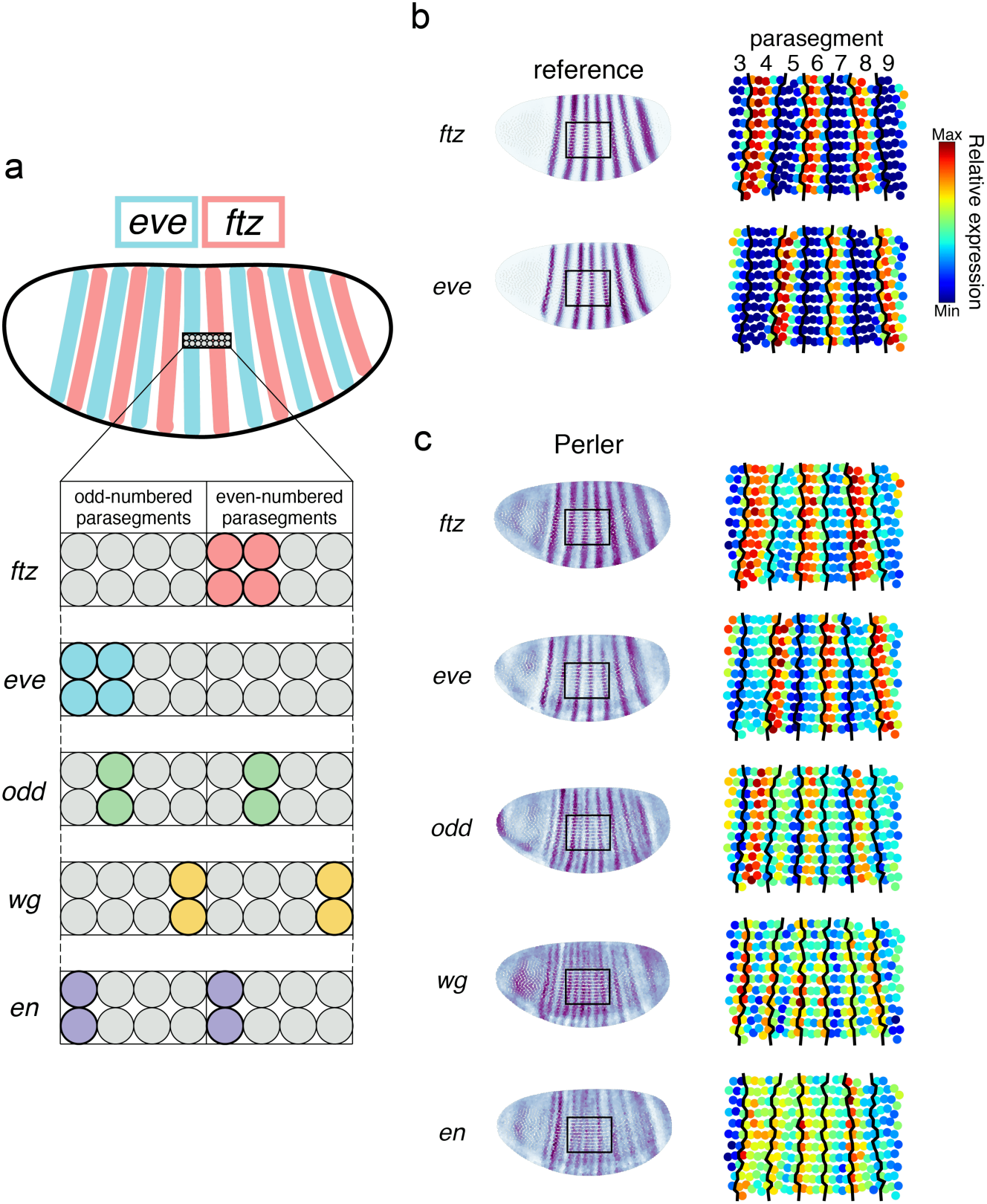
Perler prediction at single-cell resolution. (a) Spatial expression profiles of pair-rule and segment-polarity genes at the single-cell level within each parasegment^27^. The referenced stripe patterns of *ftz* and *eve*. (c) The reconstructed stripe patterns of *ftz, eve, odd, wg*, and *en* at single-cell resolution. The left panel shows the spatial gene-expression profiles generated by Perler. The right panel shows the expanded image of the left panel. The black square in the left panels is the region of interest in the right panels. (b) Black lines in the right panels indicate the boundaries of each parasegment, which were determined by expression patterns of *ftz* and *eve* in the reference ISH dataset.

In addition to the landmark genes, Perler successfully predicted the spatial expression profiles of non-landmark genes along both anterior–posterior (A–P) and dorsoventral (D–V) axes (**Fig. 3a and b**). Furthermore, we evaluated the predicted spatial profile of 308 spatially restricted genes (SRGs) proposed by Bageritz *et al*.^21^ (**Supplementary Figs. 4 and 5**) and found that Perler was able to uncover the unknown spatial gene-expression pattern. Notably, we observed that spatial patterns predicted by Seurat (v.3), Liger, and DistMap were incomplete. For example, the predicted stripes disappeared in the ventral part of embryos (e.g., *abd-A* and *Ubx* in **Fig. 3a**), whereas this issue was not observed with Perler, which accurately predicted the stripe pattern, even in the ventral part of embryos.

**Figure 5:**
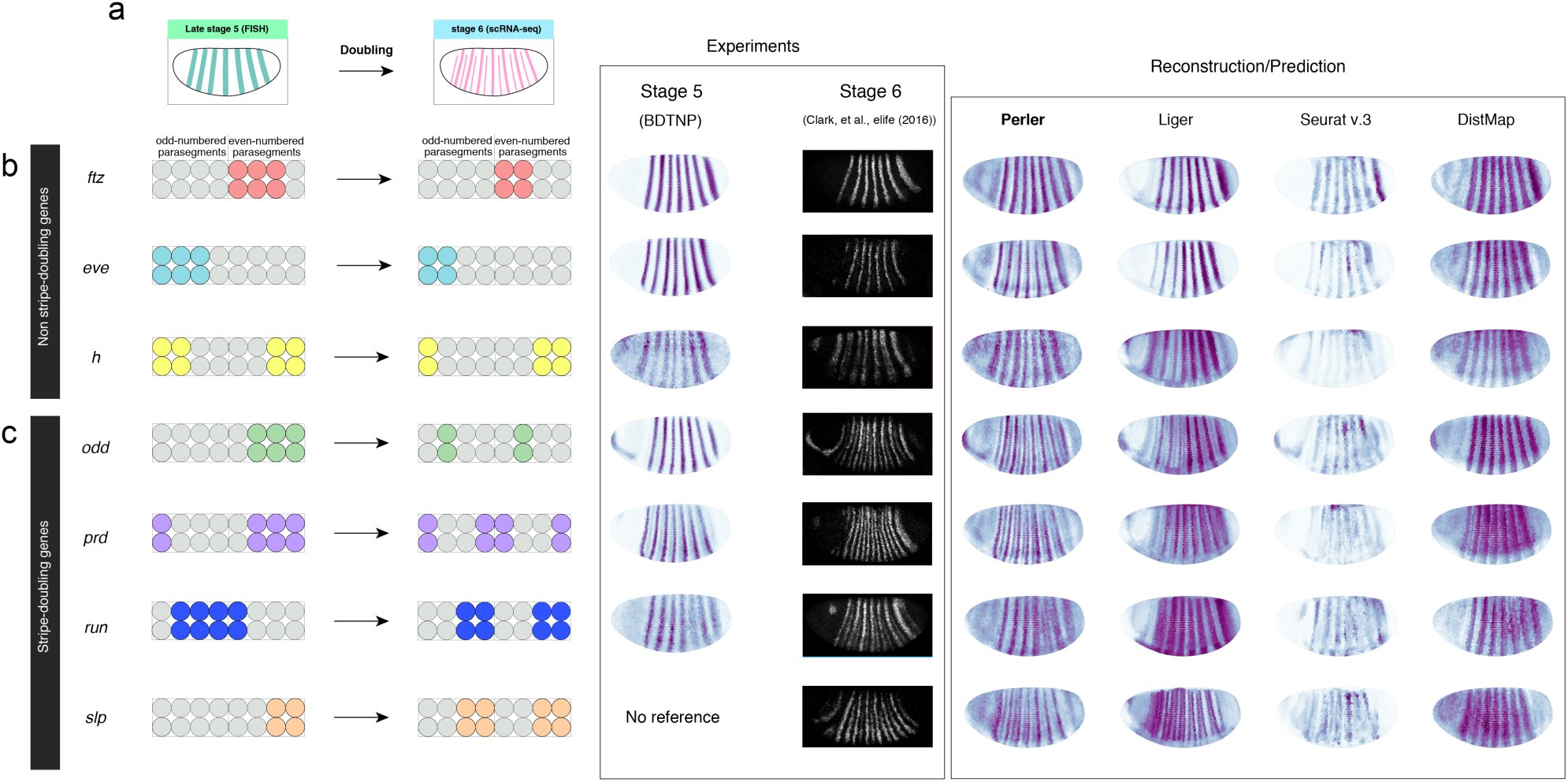
Generalization to spatial reference maps by Perler. (a) Stripe-doubling of pair-rule genes from *Drosophila* developmental stage 5 (FISH experiment) to stage 6 (scRNA-seq experiment). (b, c) Left panels: images of expression changes in (b) non-stripe-doubling genes and (c) stripe-doubling genes. Middle panels: experimental ISH data of stage 5 (BDTNP) and stage 6 (Clark et al.^27^). Right panels: gene expression reconstructed/predicted by Perler, Liger, Seurat (v.3), and DistMap. Note that *slp* expression was predicted. Experimental ISH data were reprinted from Clark et al.^26^ under a Creative Commons License (Attribution 4.0 International License).

### Prediction of 14-stripe patterns of segment-polarity genes

We then presented the spatial predictions of ‘segment-polarity’ genes, which are expressed in a 14-stripe pattern consistent with the parasegments that subdivide the trunk (main body) region of embryos (**Fig. 3b**)^22–26^. Although the BDTNP reference does not contain information concerning the genes expressed in the 14-stripe pattern, we found that Perler accurately predicted the spatial expression patterns of these segment-polarity genes, including *engrailed* (*en*), *wingless* (*wg*), *hedgehog* (*hh*), and *midline* (*mid*) (**Fig. 3b**)^22–26^. By contrast, all of the previous methods exhibited issues regarding prediction of the 14-stripe patterns. The predicted patterns demonstrated that DistMap and Seurat (v.3) were unable to predict any 14-stripe patterns, and that Liger partially predicted 14-stripe patterns, although the ventral part of each stripe was missing (**Fig. 3b**). These results suggested that Perler more accurately revealed the spatial gene-expression patterns of non-landmark genes.

We further analyzed the details of the gene-expression profiles of the segment-polarity genes within each parasegment. Each parasegment shows a four-cell width and is delimited by periodic expression of pair-rule genes and segment-polarity genes at the single-cell width resolution at stage 6^27,28^ (**Fig. 4a**). First, we confirmed that the reconstructed patterns of *ftz, eve*, and *odd* were consistent with experimental results (**Fig. 4b**). Additionally, the predicted stripes of *wg* were identified adjacent to the predicted stripes of *en*, and the predicted stripes of *en* were identified adjacent to the reconstructed stripes of *odd* (**Fig. 4b and c**). These results were consistent with experimental results^27,28^, strongly supporting the ability of Perler to reveal differences in spatial gene expression at single-cell resolution.

### Preservation of timing information of scRNA-seq data

We then investigated the effect of timing differences between scRNA-seq (stage 6) and FISH (stage 5) experiments. Although most gene-expression patterns at stage 6 are the same as those at stage 5, several “pair-rule” genes (*odd, prd, slp1*, and *run*) exhibit stripe-doubling from the 7- to the 14-stripe expression patterns during stages 5 and 6^27^ (**Fig. 5a**). Accordingly, the scRNA-seq data should intrinsically contain information for the 14-stripe expression pattern. Therefore, we determined whether Perler could reconstruct the 14-stripe pattern from the stage 6 scRNA-seq data.

In our reconstruction, *ftz, eve*, and *h* showed a 7-stripe pattern, which was consistent with the previous report^26^ showing that these genes do not exhibit stripe-doubling during stages 5 and 6 (**Fig. 5b**). For *odd, prd*, and *slp1*, which exhibit stripe-doubling, Perler reconstructions resulted in 14-stripe patterns (**Fig. 5c**). Additionally, reconstruction of *run* resulted in a partial stripe-doubling pattern, where the third stripe from the posterior of the embryo was split into two stripes (**Fig. 5c**), surprisingly suggesting that Perler detected the ongoing phase of a 7-stripe to 14-stripe pattern. These results showed that Perler was able to reconstruct embryos according to the timing of the scRNA-seq experiment. By contrast, Seurat (v.3) and DistMap reconstructed every pair-rule gene as 7-stripe patterns (**Fig. 5b and c**). Moreover, Liger reconstructed *odd, prd*, and *slp1* as broad primary seven stripes with weak secondary seven stripes, which were so obscure that it was difficult to distinguish 14 stripes, and reconstructed *run* as a 7-stripe pattern (**Fig. 5c**). These results indicated that previous methods reconstructed embryos according to the timing of FISH experiments rather than that of scRNAseq experiments. Taken together, these findings showed that Perler successfully reconstructed spatial gene-expression profiles according to the timing of scRNA-seq experiments (stage 6), regardless of the timing of FISH experiments (stage 5), while all other methods reconstructed those at the timing of FISH experiments. We concluded that Perler has the ability to not over-fit to ISH data and robustly preserve timing information in scRNA-seq data.

### Application to other datasets

To evaluate Perler applicability to other datasets, we evaluated it using three published datasets. First, we applied Perler to the zebrafish embryo datasets, in which the spatial reference map was binarized based on traditional measurement by ISH^4^ (**Fig. 6a**; see Methods). Cross-validation demonstrated that Perler accurately predicted spatial gene-expression profiles compatible with Seurat (v.1)^4^ (median receiver operating characteristic (ROC) score = 0.97), even using the binary spatial reference map (**Fig. 6a–c**).

**Figure 6:**
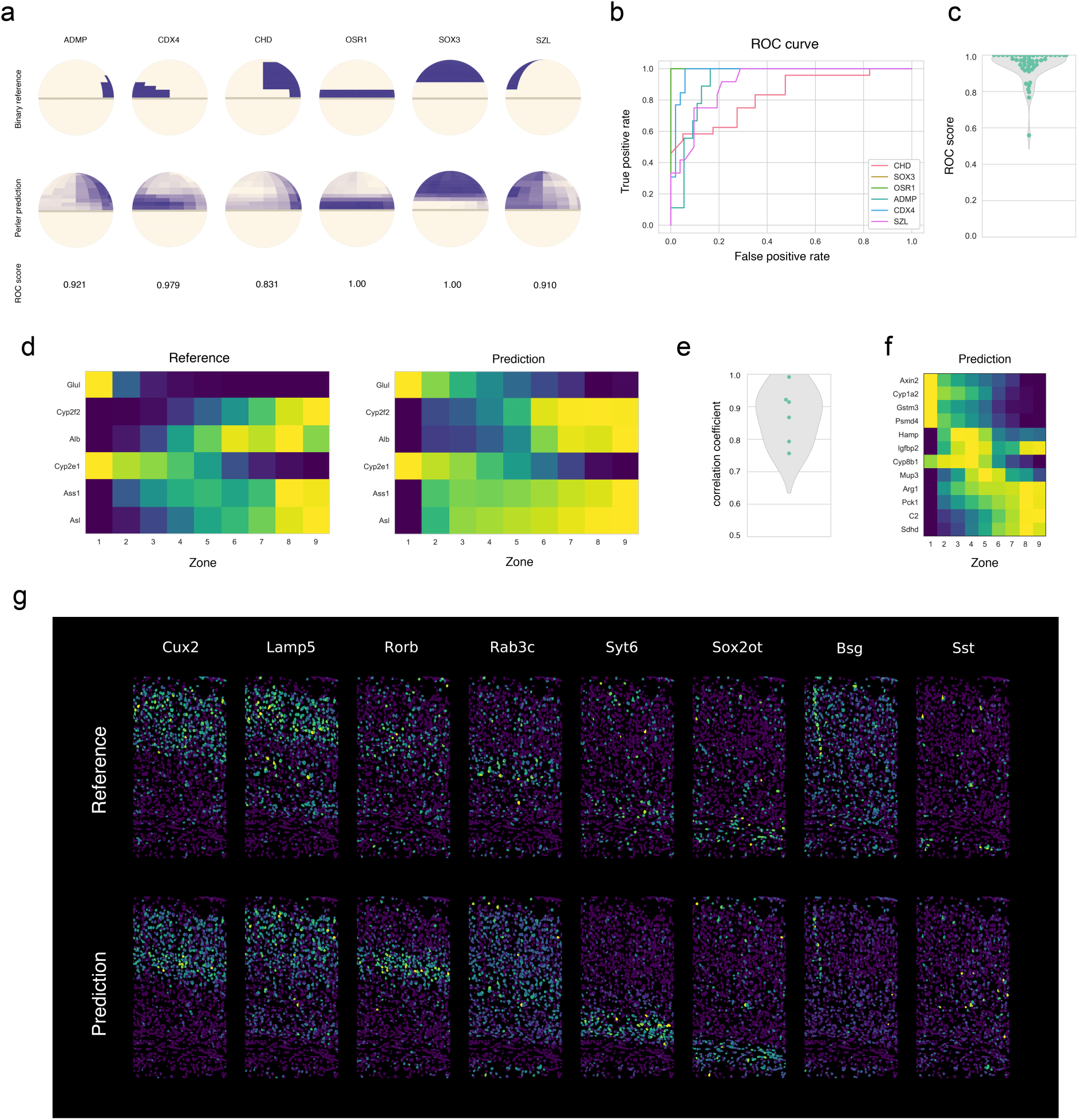
Applications of PERLER to other data. (a–c) Application of Perler for early zebrafish embryo data. (a) LOOCV experiments. The upper and lower panels show the referenced ISH data and predicted gene-expression profiles, respectively. (b) ROC curve for the LOOCV experiments for the genes shown in (a). (c) Violin plot for the predictive accuracies of Perler for the LOOCV experiments for all genes in the reference ISH data according to ROC score. (d–f) Application of Perler for mammalian liver lobules. (d) LOOCV experiments. The left and right panels show the reference ISH data and predicted gene-expression profiles, respectively. All genes from the ISH data are shown. (e) Violin plot for the predictive accuracies of Perler for the LOOCV experiments for all genes in the reference ISH data. (f) Prediction of non-landmark genes. In addition to the monotonic gene-expression profiles, non-monotonic gene-expression profiles are observed (*Hamp, Igfbp2, Cyp8b1*, and *Mup3*)^7^. (g) Application of Perler for the mouse visual cortex. The upper and lower panels show the referenced ISH data and the predicted gene-expression profiles, respectively.

We then applied Perler to mammalian liver datasets, in which the spatial reference map was measured by single-molecule (sm)FISH^7^ (**Fig. 6d**, see Methods). Cross-validation showed that the predictive accuracy (aCC = 0.87) was sufficiently high (**Fig. 6e**), and that Perler successfully predicted both monotonic and non-monotonic gene-expression gradients (**Fig. 6f**)^7^. Finally, we applied Perler to adult mouse visual cortex datasets, in which the single-cell resolution ISH data for 1,020 genes was measured by recent *in situ* technology (STARmap^15^), and scRNA-seq data for 14,739 cells available from the Allen Brain Atlas^16^. Cross-validation revealed that Perler predicted the spatial expression patterns of genes according to both layer-specific expression and cell-type-specific expression in brain cortex (**Fig. 6g**). These results suggested that Perler is applicable to prediction using high-dimensional spatial reference maps.

Taken together, the findings support Perler as a powerful tool for predicting spatial gene-expression profiles in any multicellular system with general applicability to any type of ISH data (e.g., binary or continuous, low to high dimension, and single-cell to tissue-level resolution).

## Discussion

In this study, we developed a model-based computational method (Perler) that predicts genome-wide spatial transcriptomes. Perler sequentially conducted a two-step computation, with the first step mapping ISH data points to the scRNA-seq space according to the generative linear model by EM algorithm (**Fig. 1b**), and the second step optimizing the weighting function used to predict spatial transcriptomes according to weighted scRNA-seq data points (**Fig. 1c and d**). Using a dataset for early *Drosophila* embryos, we demonstrated that Perler accurately reconstructed and predicted genome-wide spatial transcriptomes and was able to robustly preserve the timing information in scRNA-seq data. (**Figs. 2–5**). Moreover, we showed that in any multicellular system, Perler displayed broad applicability to any type of ISH data (**Fig. 6**).

### Difference from existing methods

Several studies have analyzed spatial gene-expression profiles from scRNA-seq data. Here, we discuss their differences from Perler (**Supplementary Table 1**). First, Perler does not require binarization of both ISH and scRNA-seq data, which differs from Seurat (v.1)^4^, the method described by Achim *et al*.^6^, and DistMap^5^, all of which binarize either or both data sets and result in loss of information related to continuous gene expression. We showed that Perler significantly improved the predictive accuracy of the *Drosophila* spatial transcriptome as compared with DistMap (**Fig. 2, Supplementary Fig. 3**). Additionally, even with data for zebrafish embryos only available in a traditional binary ISH dataset, Perler showed similar predictive performance with that of Seurat (v.1) (**Fig. 6**).

Perler does not need ISH data with a gene-expression distribution for each cell or subregion, which differs from the method of Halpern *et al*.^7^, which repeated smFISH experiments in order to sample distributions of landmark gene expression on a one-dimensional radial coordinate in hepatic lobules. Using the sampling distributions, their Bayesian method computed posterior probabilities of each scRNA-seq data point being generated from cells/subregions in the tissue. However, repeating smFISH experiments is obviously labor intensive in the case of two- or three-dimensional tissue. Perler addresses this sampling problem by estimating gene-expression distributions in each cell from the tissue sample using the generative linear model. This enables Perler to calculate posterior probabilities for two- and three-dimensional tissues (**Figs. 2c and 6**).

Perler is a model-based method, which differs from Seurat (v.3)^9^ and Liger^10^, both based on model-free mapping between ISH and scRNA-seq data (e.g., CCA and NMF methods). Their model-free mapping addresses gene expression as continuous variables with applicability to any kind of multicellular system; however, these methods freely map ISH data to scRNA-seq data without any assumptions (i.e., they do not account for latent relationships between the two datasets). Indeed, we showed that Seurat v.3 and Liger over-fit to the timing of ISH experiments by focusing on the stripe-doubling of pair-rule genes in *Drosophila* (**Fig. 5**). To guarantee generalized performance, we introduced generative linear modeling with biologically interpretable constraints and statistically reasonable distances. For the former, expression levels are linearly correlated between ISH and scRNA-seq measurements with gene-specific sensitivity, background signals, and noise intensity. For the latter, the pairwise distances between ISH and scRNAseq data points were evaluated in a variance-scaled manner using Mahalanobis’ metric of Gaussian mixture distribution (see Methods). Consequently, Perler avoided the over-fitting problem encountered by previous methods. Indeed, Perler reconstructed the stripe-doubling of pair-rule genes according to the timing of scRNA-seq data (**Fig. 5**).

It is worth mentioning a recent method called novoSpaRc^29^. This method proposed a new concept for predicting spatial expression patterns using the physical information of cells in tissue, which enables these predictions with little or no information regarding ISH gene-expression patterns. However, in practice, their predictive ability using *Drosophila* scRNA-seq data is unsatisfactory at single-cell resolution; therefore, this concept of using cellular information remains challenging. As a focus of future study, it would be interesting to extend our generative model to introduce prior knowledge of physical information.

### Application to multi-omics analysis

We demonstrated that Perler can integrate two distinct datasets of scRNA-expression profiles while also avoiding overfitting to the reference. These features suggest that Perler could be a suitable theoretical framework for integrating not only two RNA-expression datasets but also two single-cell datasets with different modalities, such as chromatin accessibility measured by a single-cell assay for transposase-accessible chromatin using sequencing and DNA methylation measured by chromatin immunoprecipitation sequencing. Particularly in terms of multi-omics analysis, where datasets from two different modalities do not exactly match and are often sampled from different individuals and using different time intervals^30,31^, Perler can potentially help integrate different types of single-cell genomics data. Thus, Perler provides a powerful and generalized framework for revealing the heterogeneity of multicellular systems.

## Methods

We developed a novel method to reconstruct spatial gene-expression profiles from an scRNA-seq dataset via comparison with a spatial reference map measured by ISH-based methods. In the spatial reference map, landmark gene-expression vectors (*D* genes; e.g., *D* = 84 in early *D. melanogaster* embryos) are available for all cells, whose locations in the tissue are known. The landmark gene-expression vector of cell *k* is represented as ***h*_*k*_**=(*h*_*k*,1_, *h*_*k*,2_,…, *h*_*k,D*_)^T^, where cells are indexed by *k* (*k* ∈ {1,2,*…, K*}), and *K* is the total number of cells in the tissue of interest. By contrast, in an scRNA-seq dataset, genome-wide expression (*D’* genes; e.g., *D’* = 8924 in early *D. melanogaster* embryos) lack information regarding cell location in tissue. The genome-wide expression vector of cell *n* is represented as ***y***_*n*_=(*y*_*n*,1_, *y*_*n*,2_,…, *y*_*n,D’*_)^T^, where cells are indexed by *n* (*n* ∈ {1,2,*…, N*}), and *N* is the total number of cells used for scRNA-seq measurement.

### Observation model

We modeled the difference between scRNA-seq and ISH measurements as

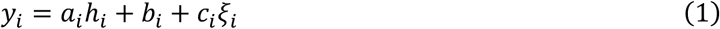

where *y*_*i*_ and *h*_*i*_ indicate expression levels of landmark gene *i* measured by scRNA-seq and ISH experiments, respectively; *ξ*_*i*_ indicates Gaussian noise with zero mean and unit variance; and *a*_*i*_, *b*_*i*_, and *c*_*i*_ are constant parameters for gene *i*, which are interpreted as scale difference amplification rates, background signals, and noise intensities, respectively.

We reduced the dimensionality of the genes to change equation (1) to

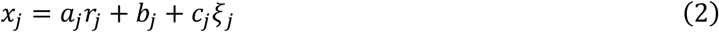

where *x*_*j*_ and *r*_*j*_ indicate expression levels of metagene *j* for scRNA-seq and ISH in the lower dimensional space, *j* ∈ {1, 2, …, *M*}; and *M* indicates the number of metagenes. In vector-matrix representation, the equation (2) is written as

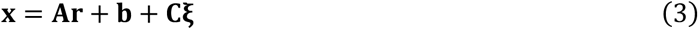

where **x** = (*x*_1_, *x*_2_, …, *x*_*Μ*_)^Τ^, **r** = (*r*_1_, *r*_2_, …, *r*_*Μ*_)^Τ^, **A** = diag(*a*_1_, *a*_2_, …, *a*_*Μ*_), **b** = (*b*_1_, *b*_2_, …, *b*_*Μ*_)^T^, **C** = diag(*c*_1_, *c*_2_, …, *c*_*Μ*_), and **ξ** =(*ξ*_1_, *ξ*_2_, …, *ξ*_*Μ*_)^Τ^.

### Metagene representation in lower dimensional space

The dimensionalities of both scRNA-seq and reference data were reduced by PLSC analysis^18^. PLSC can extract the correlated coordinates from both datasets. In PLSC analysis, the cross-correlation matrix of scRNA-seq and ISH data is first calculated as

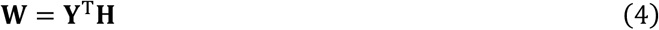

where **Y** and **H** indicate a *D*×*N* scRNA-seq data matrix with *D* landmark genes and *N* cells, and a *D*×*K* ISH data matrix with *D* landmark genes and *K* cells, respectively. **W** is then subjected to singular value decomposition as

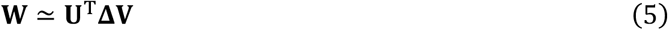

where **U, Δ**, and **V** indicate the *M×N* singular vector matrices, the *M*×*M* diagonal matrix, and *M×K* singular vector matrices, respectively, with *M* representing the reduced dimension (i.e., the number of metagenes). In this study, the metagene vectors for scRNA-seq (**x**_*n*_) and the reference data (**r**_*k*_) were respectively calculated by

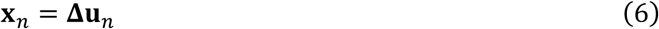

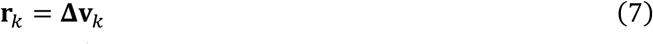

where **u**_*n*_ and **v**_*k*_ indicate the *n*^th^ row vector of **U** and the *k*^th^ row vector of **V**, respectively.

### A Gaussian mixture model (GMM) for scRNA-seq observation

We used equation (3) to transform ISH observations into scRNA-seq observations. To infer from which cells in the tissue the scRNA-seq observations originated, we developed a generative model for metagene-expression vectors for scRNA-seq data ***x***, which was expressed by a K-components GMM:

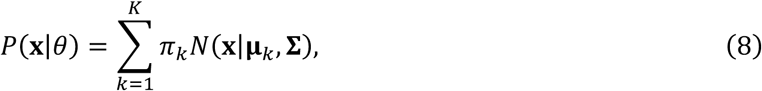

Where

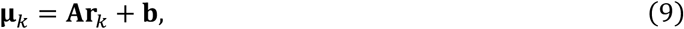

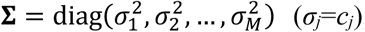, *N*(**x**|**μ, Σ**) indicates a multivariate Gaussian distribution with mean and variance-covariance matrix **Σ**, and *π*_*k*_ is the probability that **x** originated from cell *k* in the tissue. Note that **A, b**, and **Σ** are unknown parameters that need to be estimated.

The log of likelihood function of this GMM model is given by

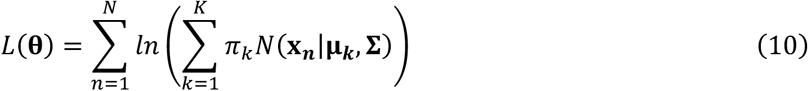

where **θ** indicates a set of the parameters **θ** ∈{**π, A, b, Σ**} and **π**=(*π*_1_, *π*_2_, …, *π*_*M*_)^T^.

### EM algorithm (the first step in Perler)

To estimate the unknown parameters (**π, A, b**, and **Σ**), we maximize the log likelihood function using the EM algorithm. In the E step, based on the current parameter values, we calculated the responsibility, which represents the posterior probability that scRNA-seq vector **x**_*n*_ was derived from cell *k* in the tissue as

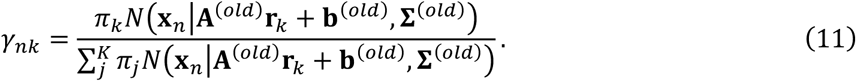

In the M step, we optimize the parameter values in order to maximize the log likelihood function based on the current responsibilities. These parameter values are updated as follows:

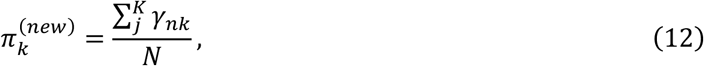

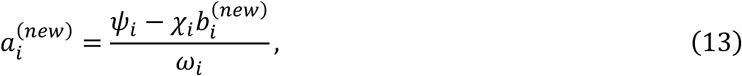

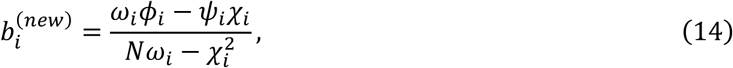

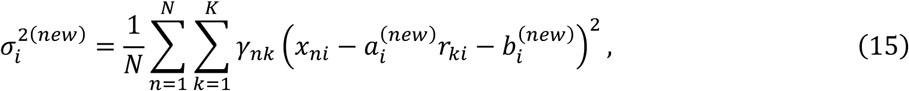

Where

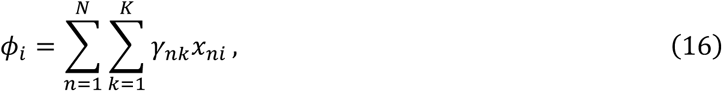

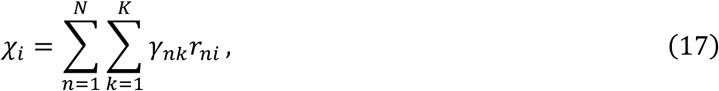

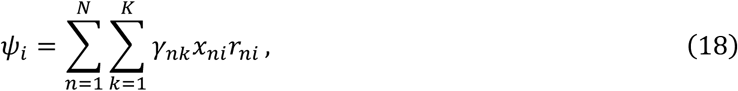

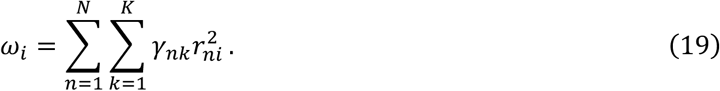

The detailed derivation for these equations is presented in a later subsection. The E and M steps iterate until the log likelihood function converges, after which the obtained estimated parameters 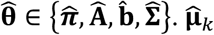 are given as

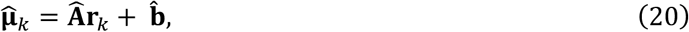

describing the mapped metagene-expression vector of cell *k* measured by ISH. Note that 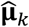 is the metagene-expression vector in the scRNA-seq space.

### Spatial reconstruction (the second step in Perler)

We reconstructed/predicted the gene-expression vector by weighted averaging all scRNA-seq data points as

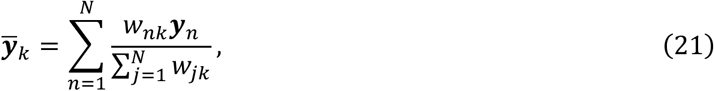

where *y*_*n*_ indicates the *n*^th^ scRNA-seq data point (*D-*component vector). *w*_*nk*_ is calculated by

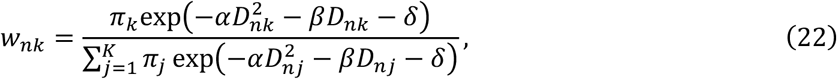

where *α, β*, and *δ* are positive constants. Note that *δ* in the numerator and denominator of equation (22) are canceled out. *D*_*nk*_ indicates Mahalanobis’ distance between scRNA-seq data point **x**_*n*_ and cell *k*:

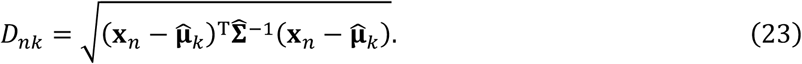

If *α* = 1/2 and *β* = 0, *w*_*nk*_ is exactly the posterior probability that scRNA-seq data point **x**_*n*_ is generated by cell *k*. Note that equation (21) has a similar structure to the Nadaraya–Watson model^13^. Values of *α* and *β* are determined by cross-validation.

### Hyperparameter optimization

We optimized the hyperparameters *α* and *β* of the weighting function by LOOCV in order to fit the predicted gene expression to the referenced gene expression measured by ISH. To this end, we removed one of the landmark genes from the ISH data and used this dataset to predict the spatial gene-expression profile of the removed landmark gene with the fixed hyperparameters in Perler. This LOO prediction was repeated for every landmark gene. We then quantitatively evaluated the predictive performance of these hyperparameters according to the mutual information existing between the predicted expression and referenced expression of all landmark genes:

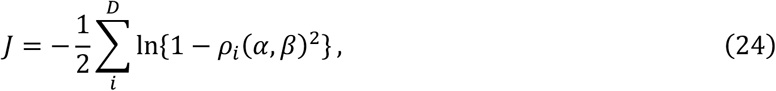

where *J* is the approximated mutual information between the predicted and referenced gene expression. *ρ*_*i*_(*α, β*) indicates the Pearson’s correlation coefficient between the predicted spatial expression pattern of each landmark gene *i* and its reference ISH data as

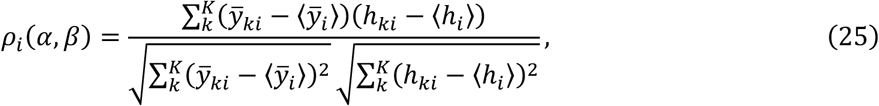

Where

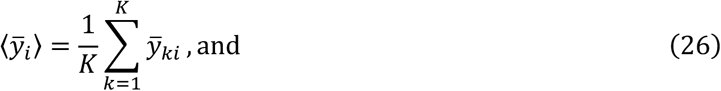

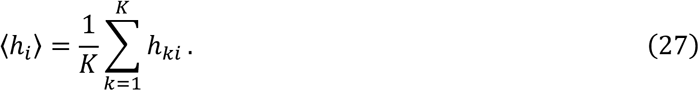

The derivation of *J* is described in a later subsection. Here, we optimized *α* and *β* by grid search in order to maximize the mutual information, *J*. We then used the optimized hyperparameters to predict the spatial profile of non-landmark genes (**Fig. 3, Supplementary Fig. 5**). To evaluate the predictive performance of Perler (**Fig. 3**), we removed each landmark gene from the mutual information and re-optimized the hyperparameters. This re-optimization is repeated for every landmark gene. Note that for the zebrafish embryo data, we used the ROC score instead of the correlation coefficient, because only the binary ISH data was available. Additionally, for the mouse visual cortex data, we conducted 10-fold cross-validation because of the massive computational cost of LOOCV for the large number of landmark genes (1,020 genes).

### Data acquisition and preprocessing

For *D. melanogaster* reconstruction, we used scRNA-seq and ISH data at *Drosophila* Virtual Expression eXplorer (DVEX) (https://shiny.mdc-berlin.de/DVEX/^5^), which was originally used for DistMap^5^. In these data sets, the number of scRNAseq data points is 1297, whereas the number of cells to be estimated in the embryos is 3039. The expressed mRNA counts in this scRNA-seq dataset were already log normalized according to the total number of unique molecular identifiers (UMIs) for each cell. For each gene, we subtracted the average expression from the scRNA-seq data. Additionally, the ISH data were log-scaled and subtracted average expression from this ISH data, as same as the scRNA-seq data.

For reconstruction of the early zebrafish embryos, we acquired the public scRNA-seq and ISH data from the Satija Lab homepage (https://satijalab.org/^4^), with these data originally used by Seurat (v.1)^4^. In these data, the number of scRNAseq data points is 851, whereas the number of subregions to be estimated in the embryos is 64. Note that the ISH data were binary. Similar to the *Drosophila* data, we log-scaled both scRNA-seq and ISH datasets and subtracted the average expression of each gene.

For reconstruction of the mammalian liver, we used scRNA-seq and smFISH data provided by Halpern *et al*.^7^. In these data, the number of scRNAseq data points is 1415, whereas the number of zones to be estimated in the embryos is 9. Because multiple samples were provided in the smFISH data, we calculated their average at each tissue location for Perler, followed by log-scaling both the scRNA-seq and smFISH data and subtracting the average expression of each gene.

For reconstruction of the mouse visual cortex, we used scRNA-seq data provided by the Allen Brain Institute^16^ and smFISH data provided by Wang *et al*.^15^, respectively, which were originally used for Seurat (v.3)^9^. The number of scRNAseq data points is 14739, whereas the number of cells to be estimated in the cortex is 1549. We log-scaled both the scRNA-seq and smFISH data and subtracted the average expression of each gene.

### Data visualization

For *D. melanogaster*, we visualized the reconstructed gene-expression profile at single-cell resolution by using the three-dimensional coordinates of all cells from DVEX (https://shiny.mdc-berlin.de/DVEX/^5^). Because the embryo is bilaterally symmetric, we mapped the reconstructed spatial gene-expression levels of the 3,039 cells in the right-half embryo. According to the previous study^5^, we then mirrored the spatial gene-expression levels of the right-half cells to the remaining cells in left-half embryo. In the case of the early zebrafish embryos, we visualized the reconstructed gene expression using the ‘zf.insitu.vec.lateral’ function of Seurat (v. ≥ 1.2)^4^. In the case of the mammalian liver, we visualized the reconstructed gene expression as a heatmap. In the case of the mouse visual cortex, we visualized the reconstructed gene expression at single-cell resolution. We used two-dimensional coordinates of all cells within cortical slices provided by Wang *et al*.^15^.

### Derivation of the EM algorithm

The goal of the EM algorithm is to maximize the likelihood function p(**X**|**θ**) with respect to ***θ***, where **X** = {**x**_1_, **x**_2_, …, **x**_*N*_} and **θ** = {**π, A, b**, Σ}. The generative model of scRNA-seq data point ***x*** with latent variables ***z*** is formulated, as follows. The probability distribution of **z** is

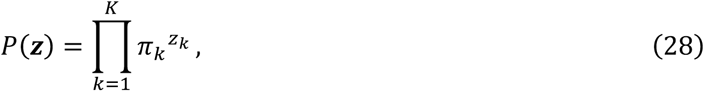

where **z** is a vector in a one-of-*K* representation that shows from which cells/regions in tissue a scRNA-seq sample originated; *z*_*k*_ is the *k*^th^ element of **z**; *K* is the number of the elements of the latent variables ***z*** equal to the number of cells in the tissue; and π_*k*_ is probability that *z*_*k*_= 1. The probability distribution of **x** conditioned by **z** is

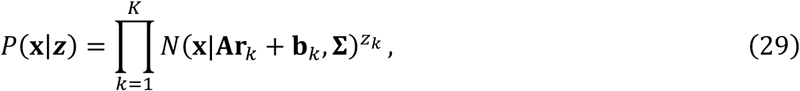

where *N*(**x**|**μ, Σ**) indicates a Gaussian distribution with mean **μ** and variance **Σ**; **A**_*k*_ is the *M*×*M* diagonal matrix; **b**_*k*_ indicates the *M* elements vector in equation (3); and **r**_*k*_ indicates the *M* elements vector describing the metagene-expression level in cell *k*. The joint probability distribution of **x** and **z** is

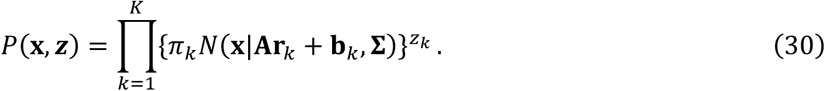

Note that the marginalized distribution of **z** becomes equation (8). The likelihood function for the complete dataset {**X, Z**} is given as

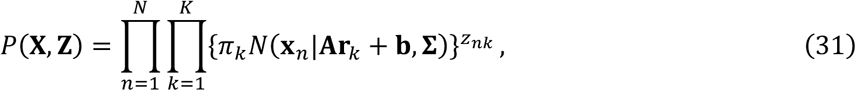

where **Z** = {**z**_1_, **z**_2_, …, **z**_*N*_}. Therefore, the expectation of its log-likelihood function over the posterior distribution of P(**Z**|**X, θ^(*old*)^**) becomes

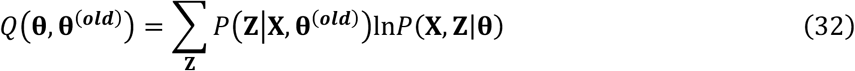

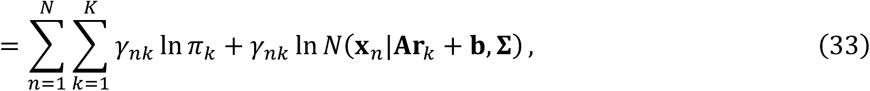

where *γ*_*nk*_ is the expectation of *z*_*nk*_ over *P*(**Z**|**X, θ^(*old*)^**) given as

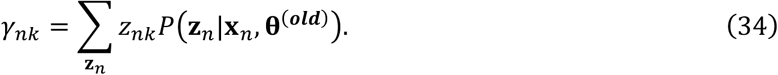

According to Bayes’ theorem,

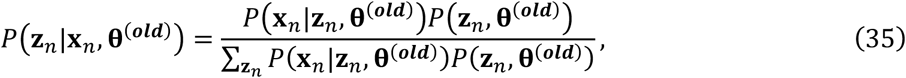

where *P*(*z*_*nk*_=1|**x**_*n*_) becomes equation (11).

In the E step, *γ*_*nk*_ is calculated based on the current parameter values of **θ^(*old*)^**. In the M step, we update the parameter values **θ** by maximizing the Q-function as

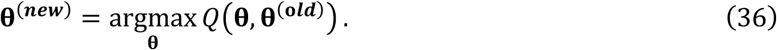

The maximization of *Q*(**θ, θ^(*old*)^**) with respective to **A, b**, and **Σ** is achieved by ∂*Q*/∂**A** = 0, ∂*Q*/∂**b** = 0, and ∂*Q*/∂**Σ**=0, leading to equations (13–19). **π** is updated by introducing a Lagrange multiplier to enforce the constraint 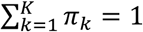, leading to equation (12).

### Derivation of mutual information

We derived equation (24) by approximating the following mutual information between the reconstructed spatial expression pattern of the landmark genes and their reference map:

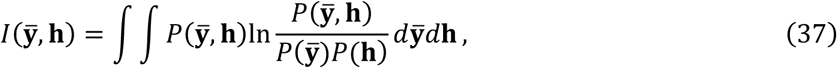

where 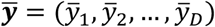, **h** = (*h*_1_, *h*_2_, …, *h*_*D*_), and 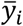 and *h*_*i*_ indicate random variables representing the predicted and referenced expression levels of landmark gene *i*, respectively, and 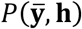 indicates the joint probability distribution of 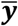 and **h**. Here, we assumed that spatial expressions of landmark genes are independent from one another, which leads to

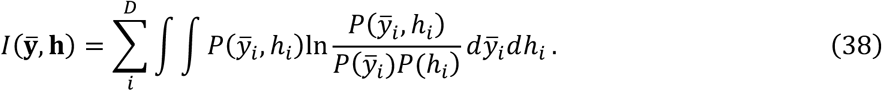

We calculated 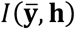 by assuming 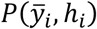 as a bivariate Gaussian distribution and obtained

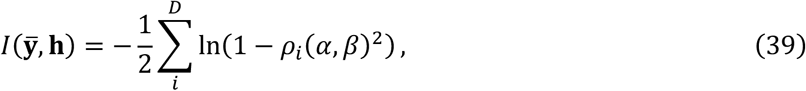

where *ρ*_*i*_(*α, β*) denotes the calculated Pearson’s correlation coefficient calculated.

## Acknowledgements

This study was supported in part by the Cooperative Study Program of Exploratory Research Center on Life and Living Systems (ExCELLS) (program Nos.18-201, 19-102, and 19-202 to H.N.); a Grant-in-Aid for Young Scientists (B) (16K16147 and 19H04776 to H.N.), a Grant-in-Aid for Scientific Research (B) (17KT0021 to T.K.) and a JSPS research fellowship for young scientist (to S.S.) from the Japan Society for the Promotion of Science (JSPS); the Naito Foundation (to T.K.); and the Keihanshin Consortium for Fostering the Next Generation of Global Leaders in Research (K-CONNEX) established by the program of Building of Consortia for the Development of Human Resources in Science and Technology, MEXT (to T.K.).

## Author Contributions

H.N., S.S. and T.K. conceived the project. Y.O., H.N. and K.N. developed the method, Y.O. implemented the software, and Y.O. and S.S. analyzed data. Y.O. and H.N. wrote the manuscript with input from all authors.

## Competing Interests

The authors declare no competing interests.

## Notes

### Competing Interest Statement

The authors have declared no competing interest.

