## Supplementary figures 1-4 for "Model-based prediction of spatial gene expression via generative linear mapping"

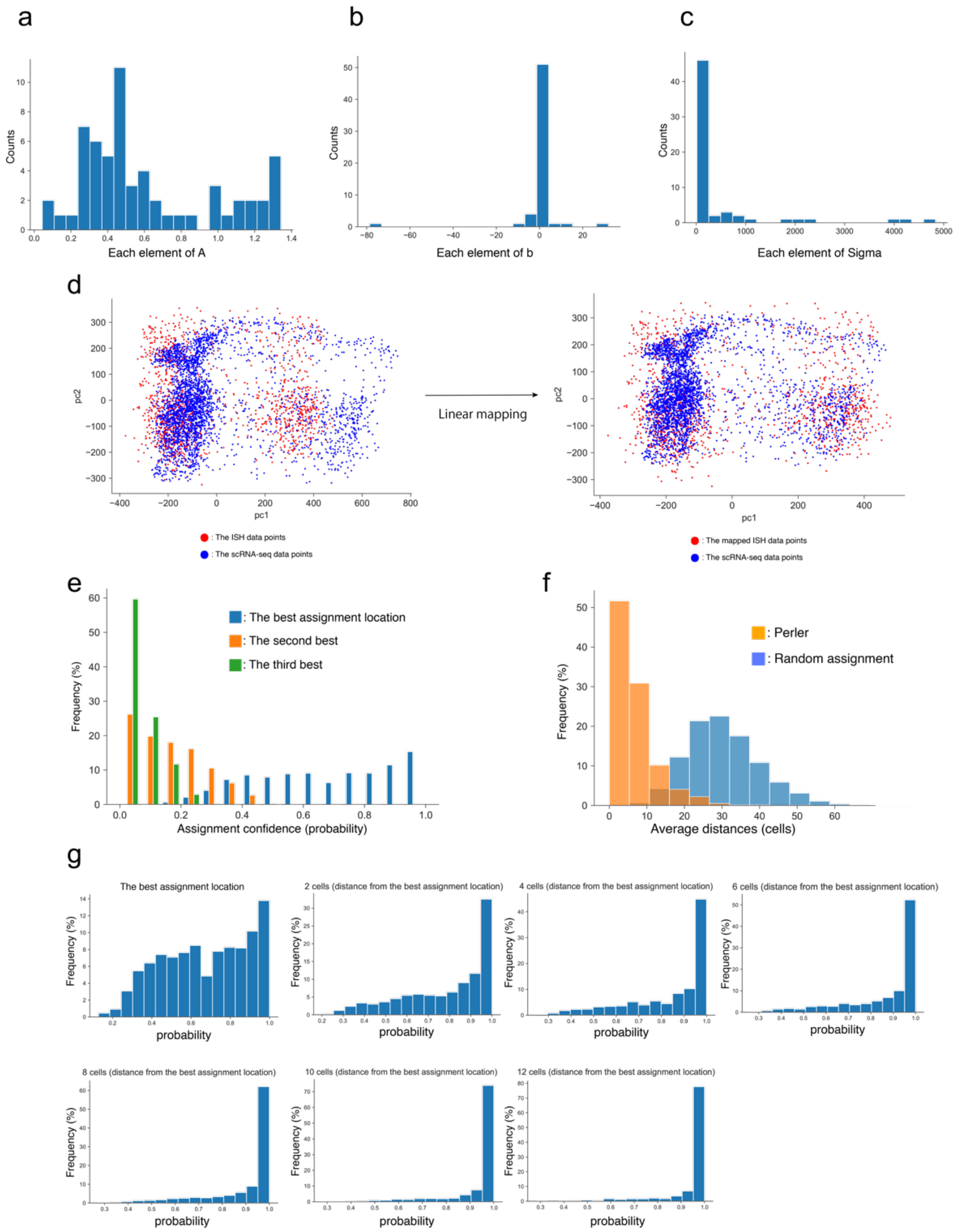


**Supplementary Figure 1: Linear mapping property of Perler**

(a–c) Histograms of the distributions of the estimated parameters of generative linear mapping: A (left), b (middle), and Σ (right) (see Methods). Note that because A and Σ are diagonal matrices, only the diagonal elements of A and Σ are shown in the middle and right panels. (d) Scatter plot of scRNA-seq and ISH data points before (left) and after (right) mapping and corresponding to Fig. 1b and Fig. 2a. Principal component analysis[13] was used to visualize high-dimensional gene-expression data into two dimensions. (e) Histogram of the posterior probabilities of the top three cells for each scRNA-seq data point. (f) Histogram of the assigned specificity evaluated by the distance between the best assigned location and the following best three locations. The distance was calculated by mean path length on the k-NN graph comprising all cells in the tissue (k = 6). (g) Histograms of the assigned confidence corresponding to Fig. 2d. Each histogram shows the detailed distributions of each boxplot in Fig. 2d.

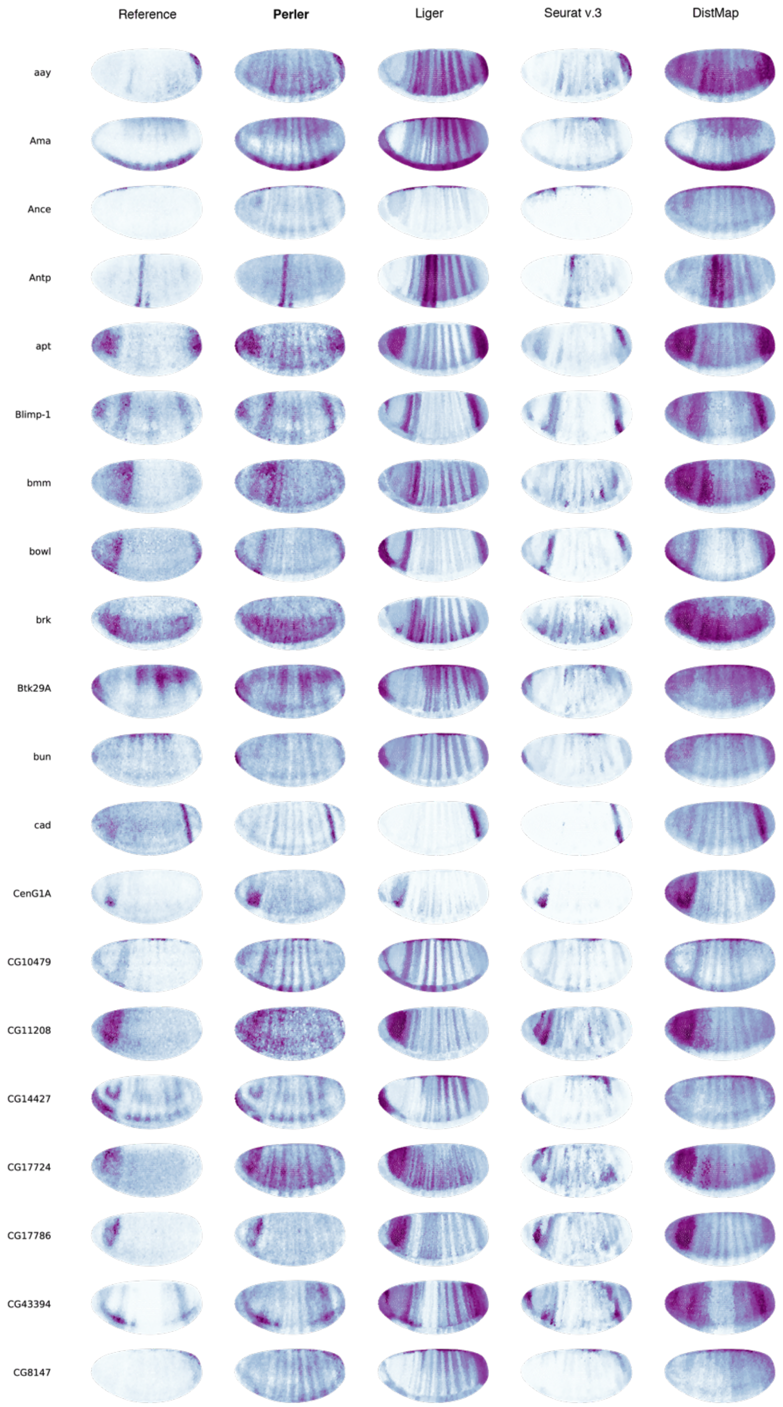


**Supplementary Figure 2: Spatial reconstruction of all landmark genes (1)**

Spatial reconstruction of all landmark genes (84 genes) by Perler, Liger, Seurat (v.3), and DistMap.

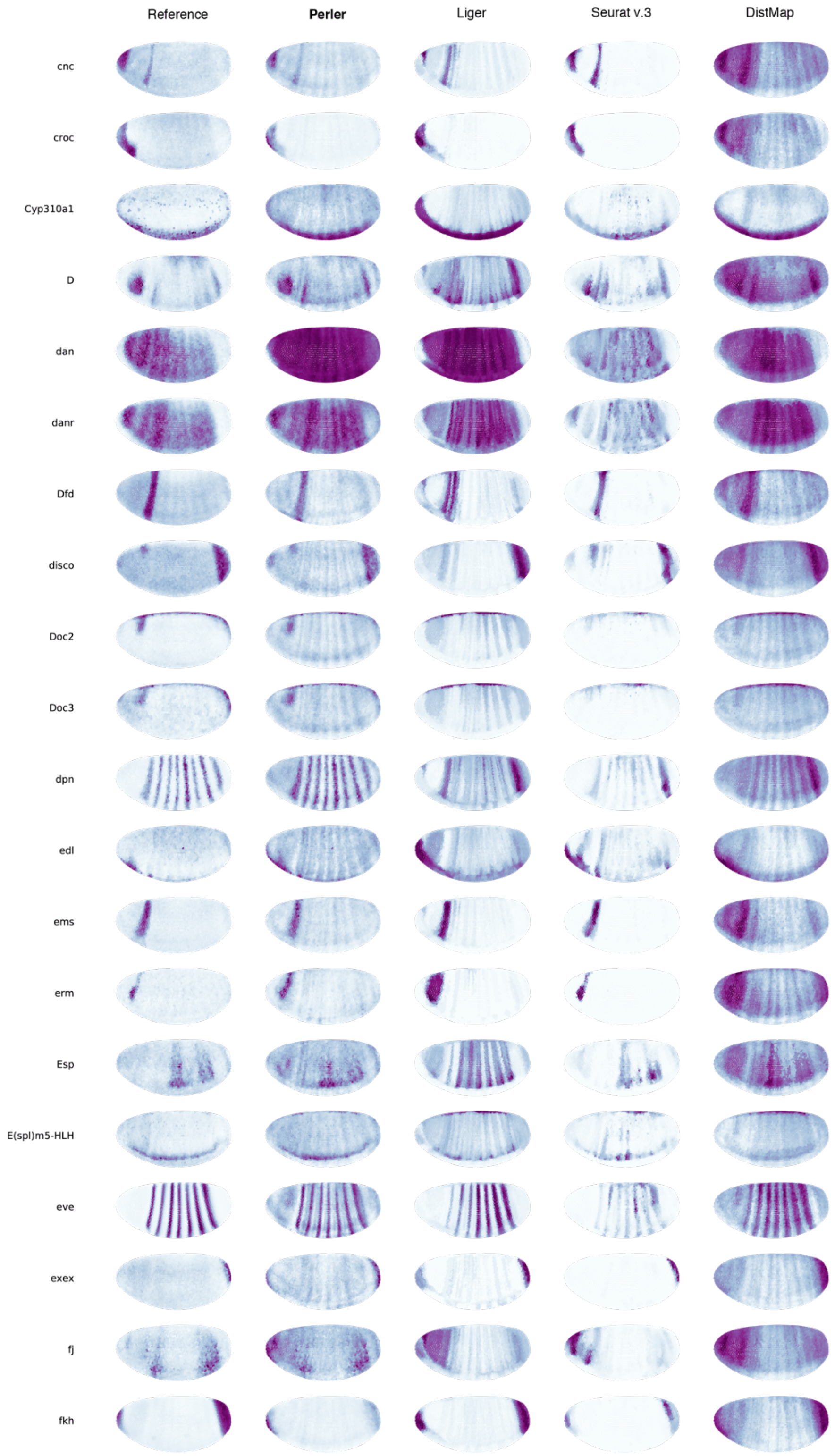


**Supplementary Figure 2 (2)**

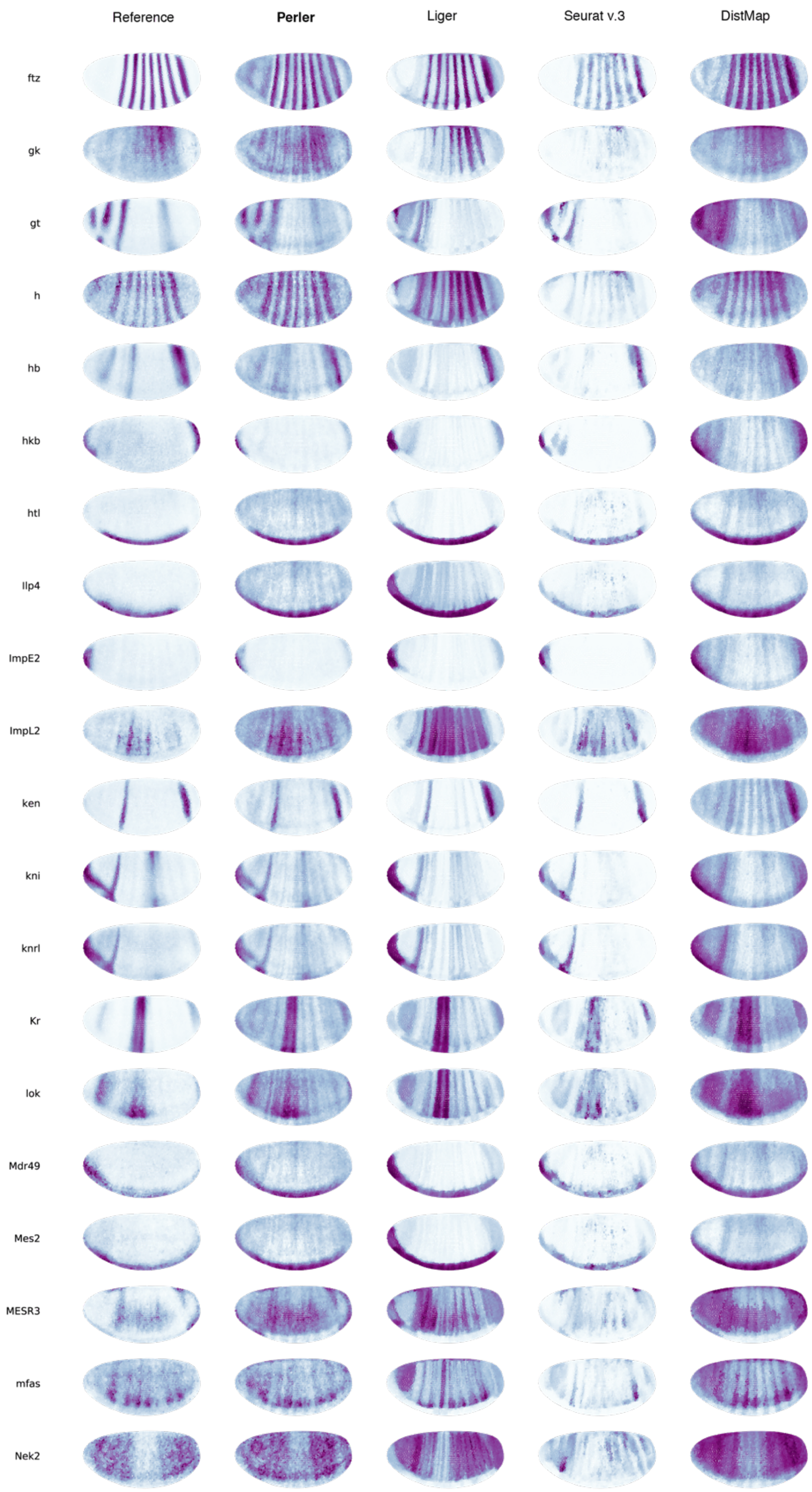


**Supplementary Figure 2 (3)**

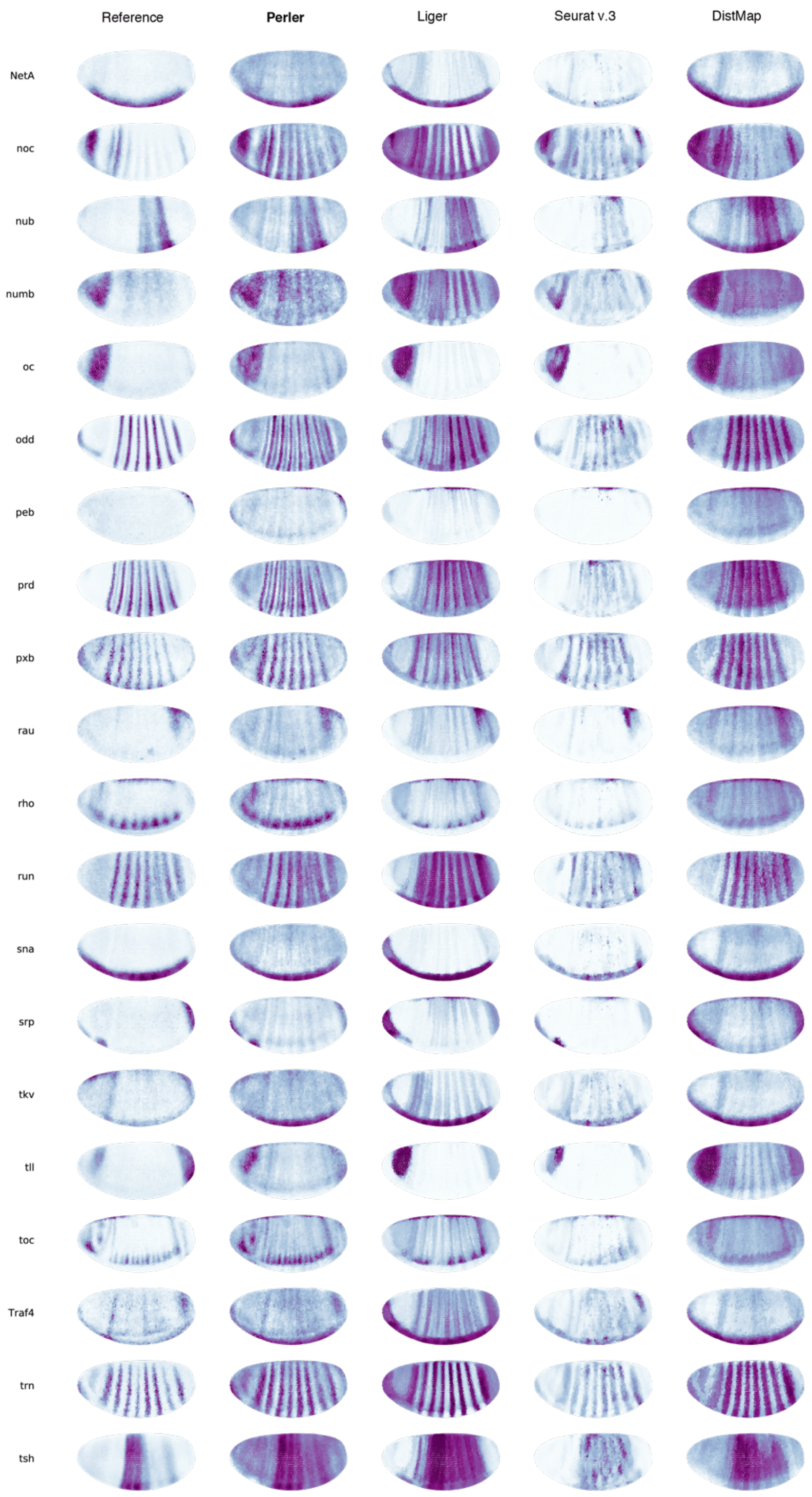


**Supplementary Figure 2 (4)**

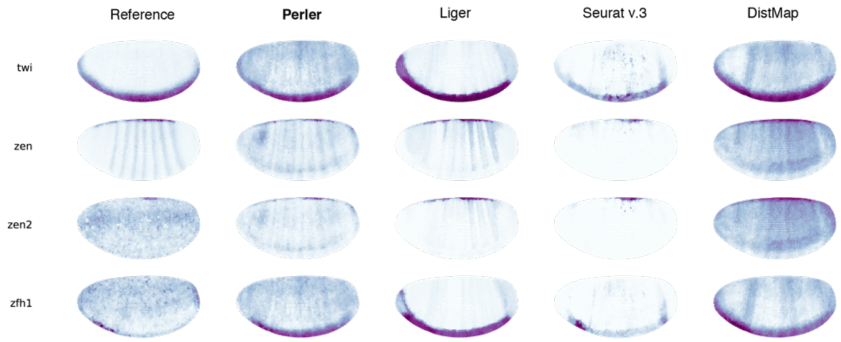


**Supplementary Figure 2 (5)**

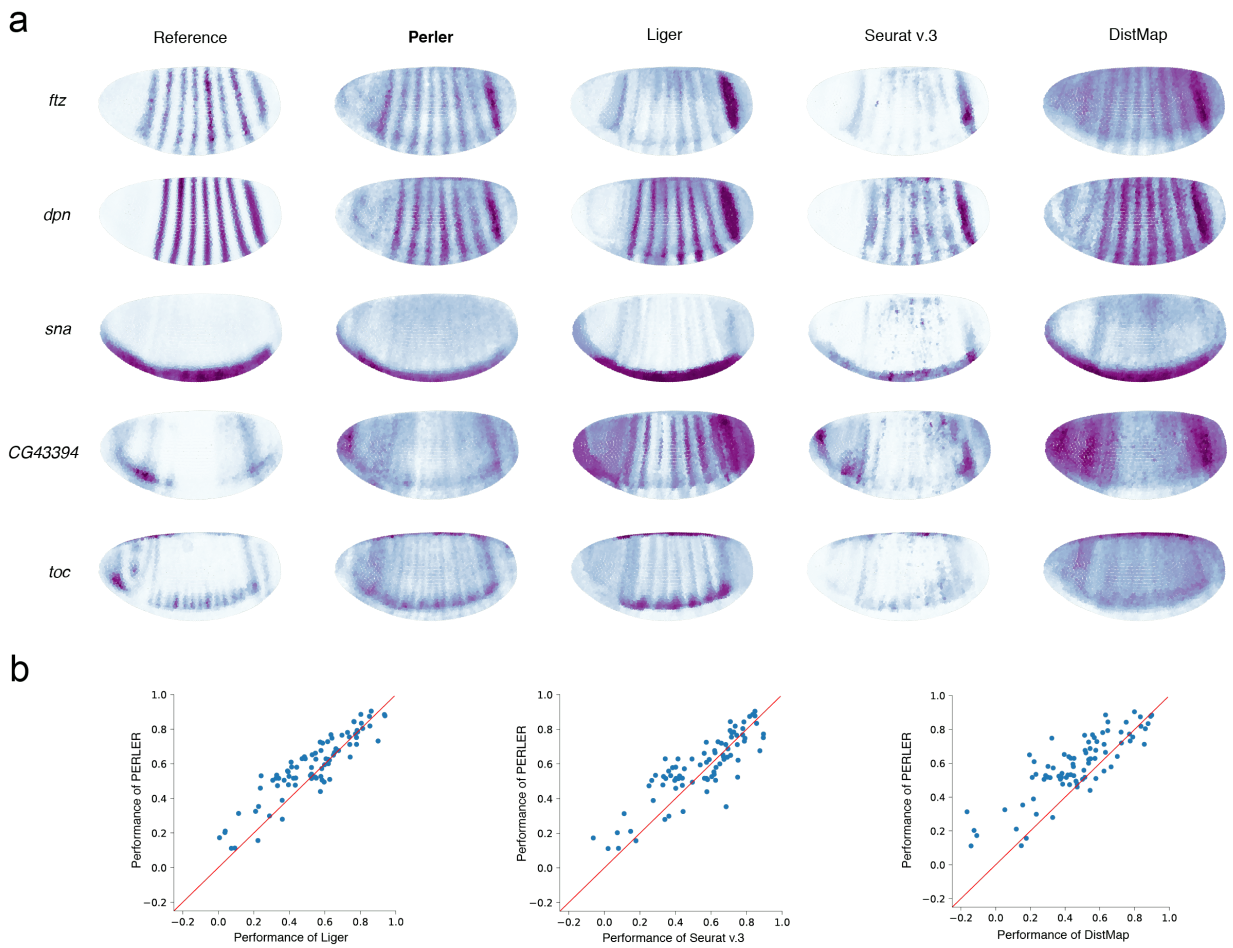


**Supplementary Figure 3: Spatial prediction of non-landmark spatially restricted genes**
(a) Predictions of landmark gene expression by Perler. Left and right panels depict the spatial reference maps and the predicted spatial gene-expression profiles. For each prediction, the predicted gene was removed from the reference ISH data (LOOCV). (b) Performance comparison of Perler with Liger (left, two-sided Wilcoxon test: $p = 2.8\times10^{-6}$), Seurat (v.3) (middle, two-sided Wilcoxon test: $p = 4.0\times10^{-3}$), and DistMap (right, two-sided Wilcoxon test: $p = 1.3\times10^{-10}$). Each dot indicates the predictive accuracies for each gene by Perler and previous methods.

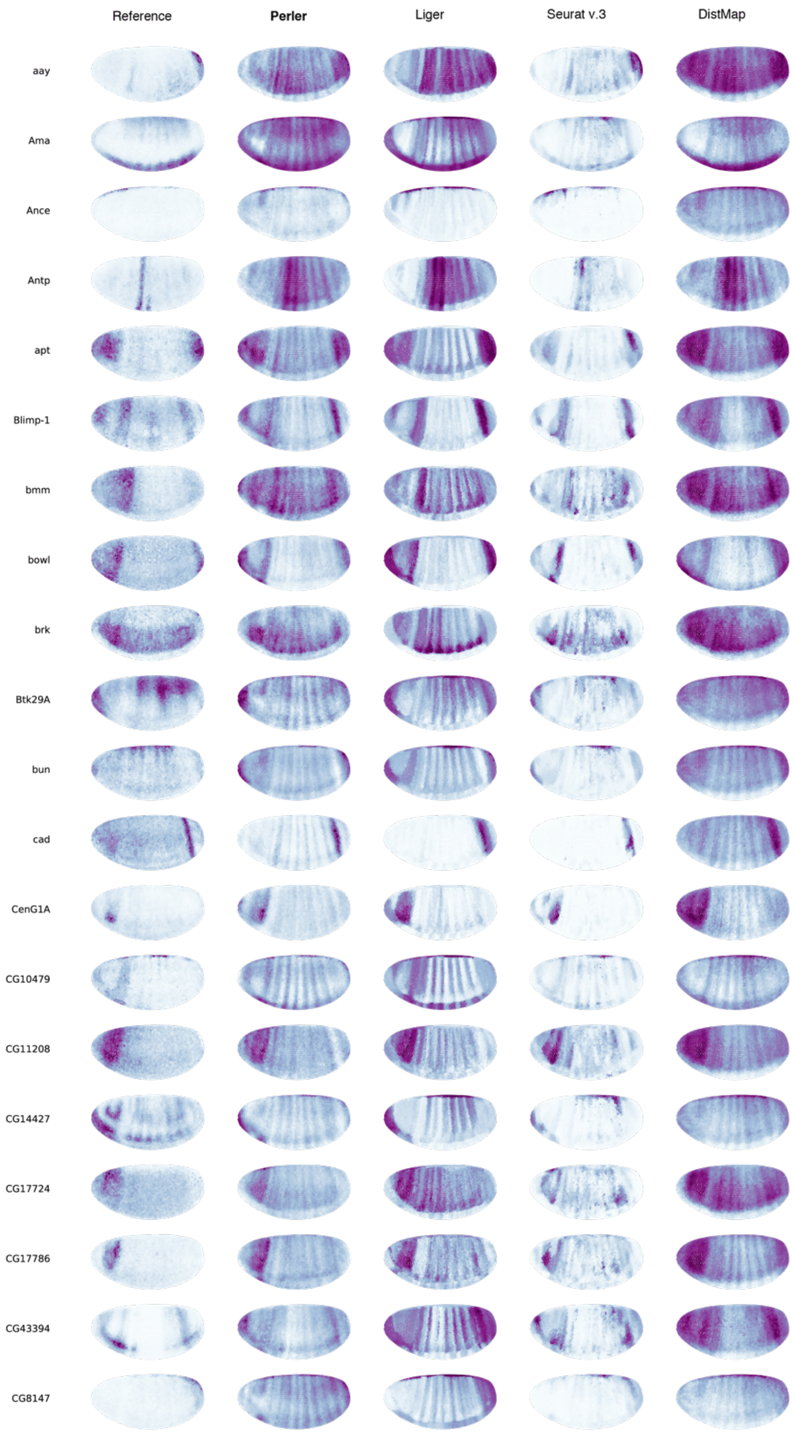


**Supplementary Figure 4: Spatial prediction of all landmark genes (1)**
Spatial prediction of all landmark genes (84 genes) by Perler, Liger, Seurat (v.3), and DistMap. The spatial prediction was generated by LOOCV experiments.

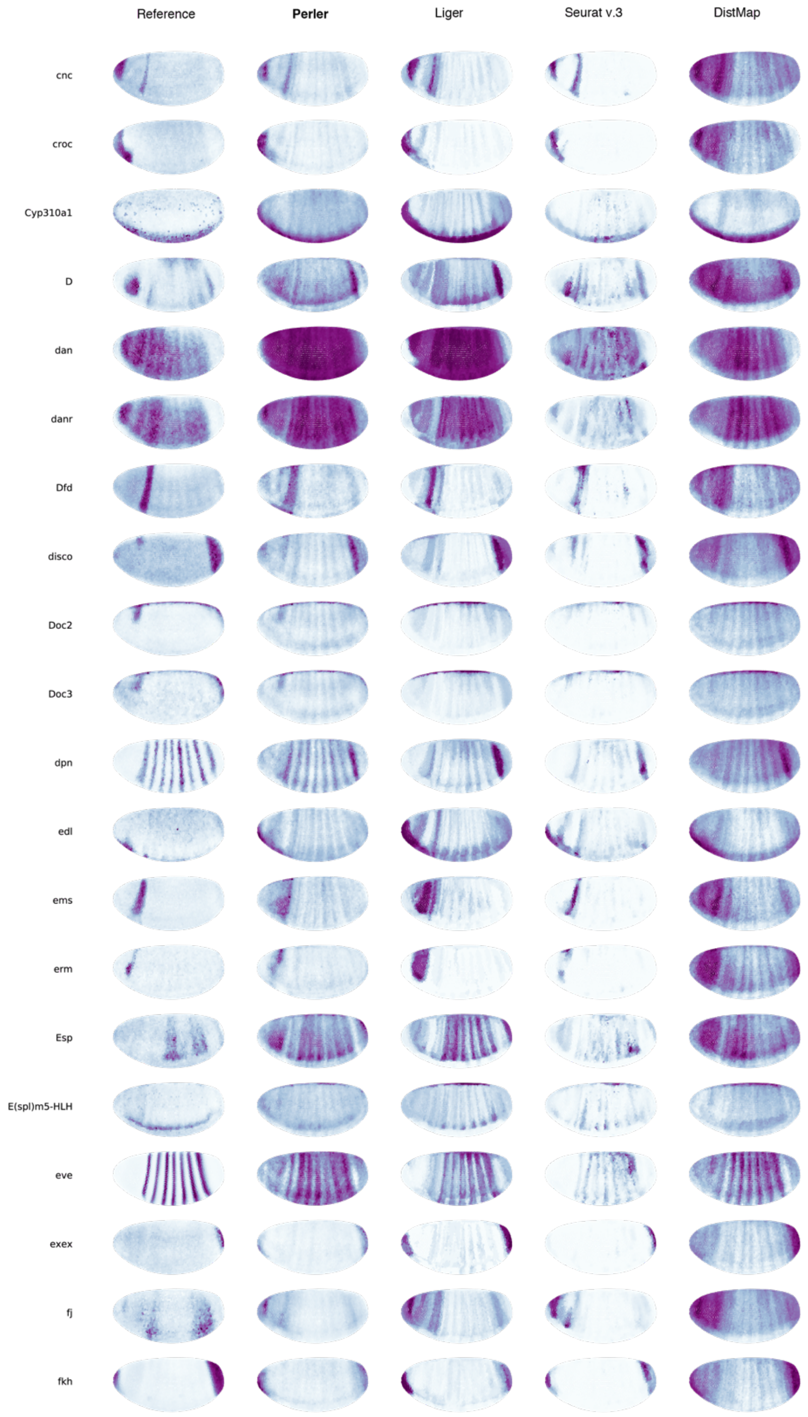


**Supplementary Figure 4 (2)**

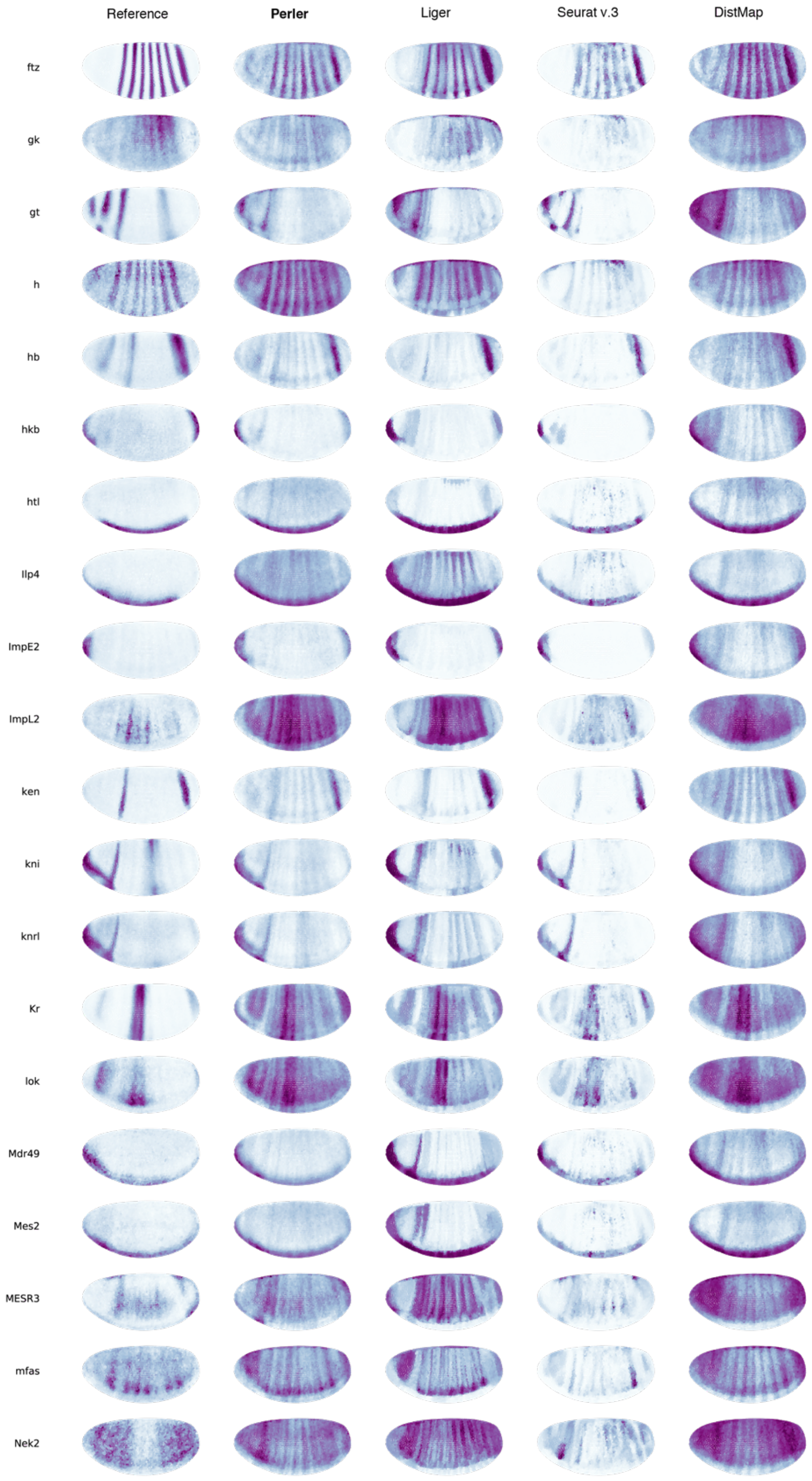


**Supplementary Figure 4 (3)**

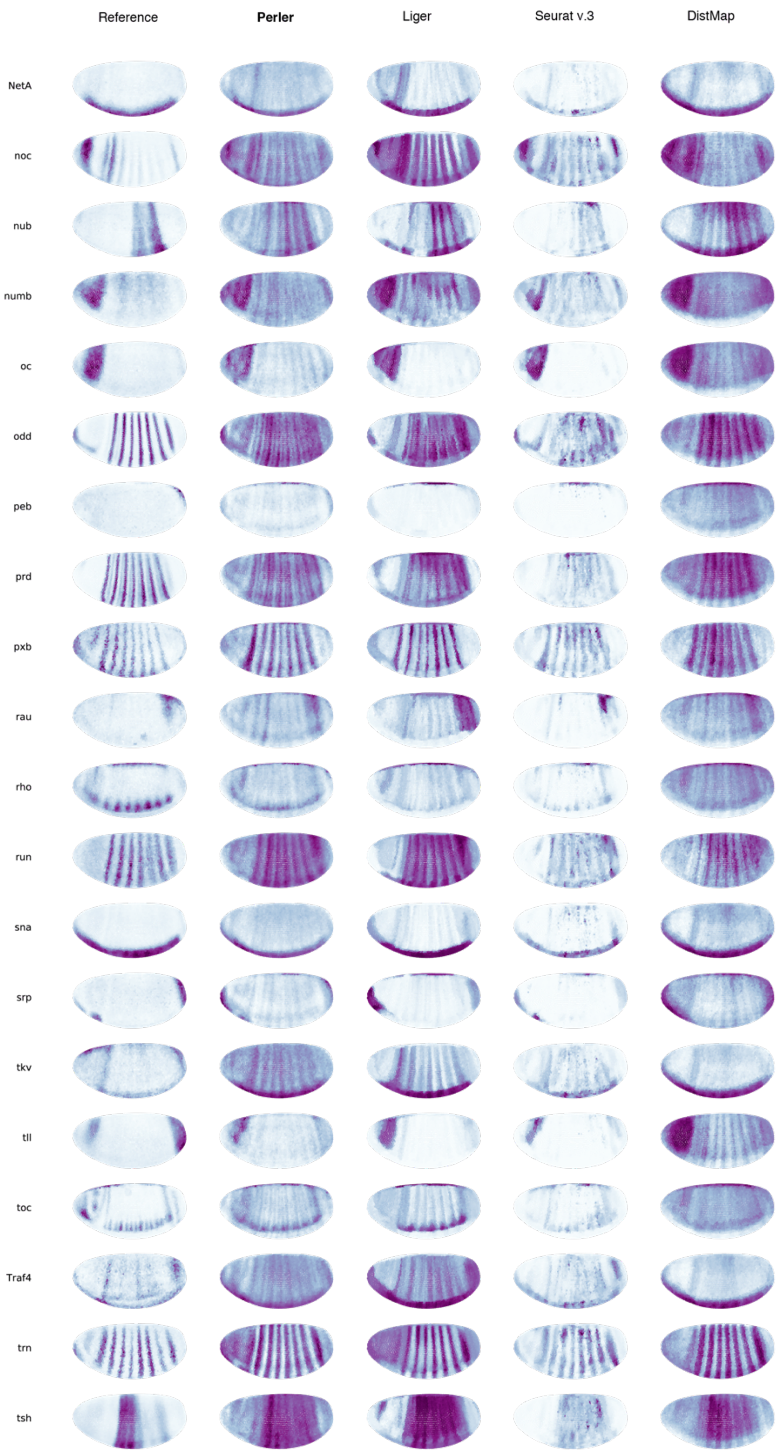


**Supplementary Figure 4 (4)**

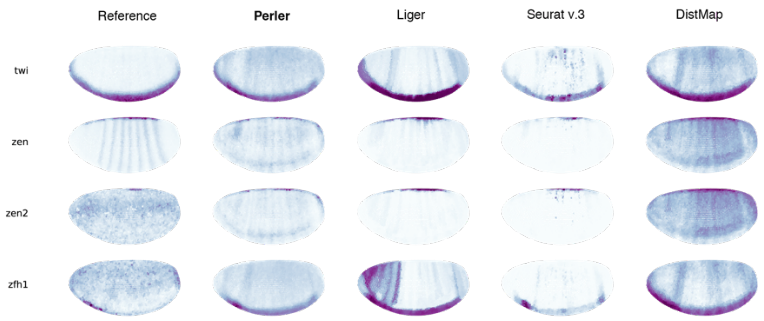


**Supplementary Figure 4 (5)**
