## Supplementary figures 5-6, table 1 for "Model-based prediction of spatial gene expression via generative linear mapping"

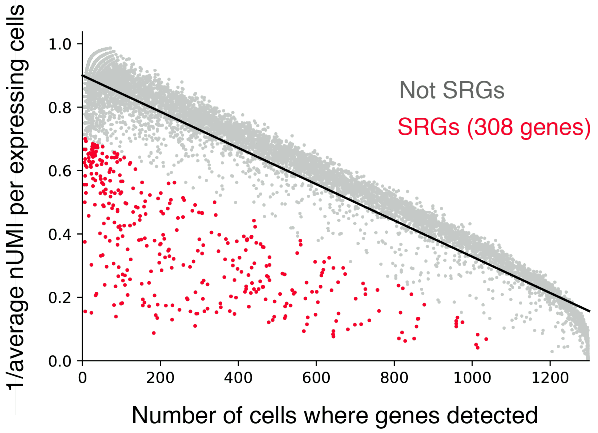


**Supplementary Figure 5: Identification of spatially restricted genes (SRGs)**
Scatter plot identifying SRGs. The red and grey points indicate SRGs and other genes, respectively. The black line indicates the linear regression. The region of SRGs was defined by the area under the minus-2 standard deviations of the regression line.

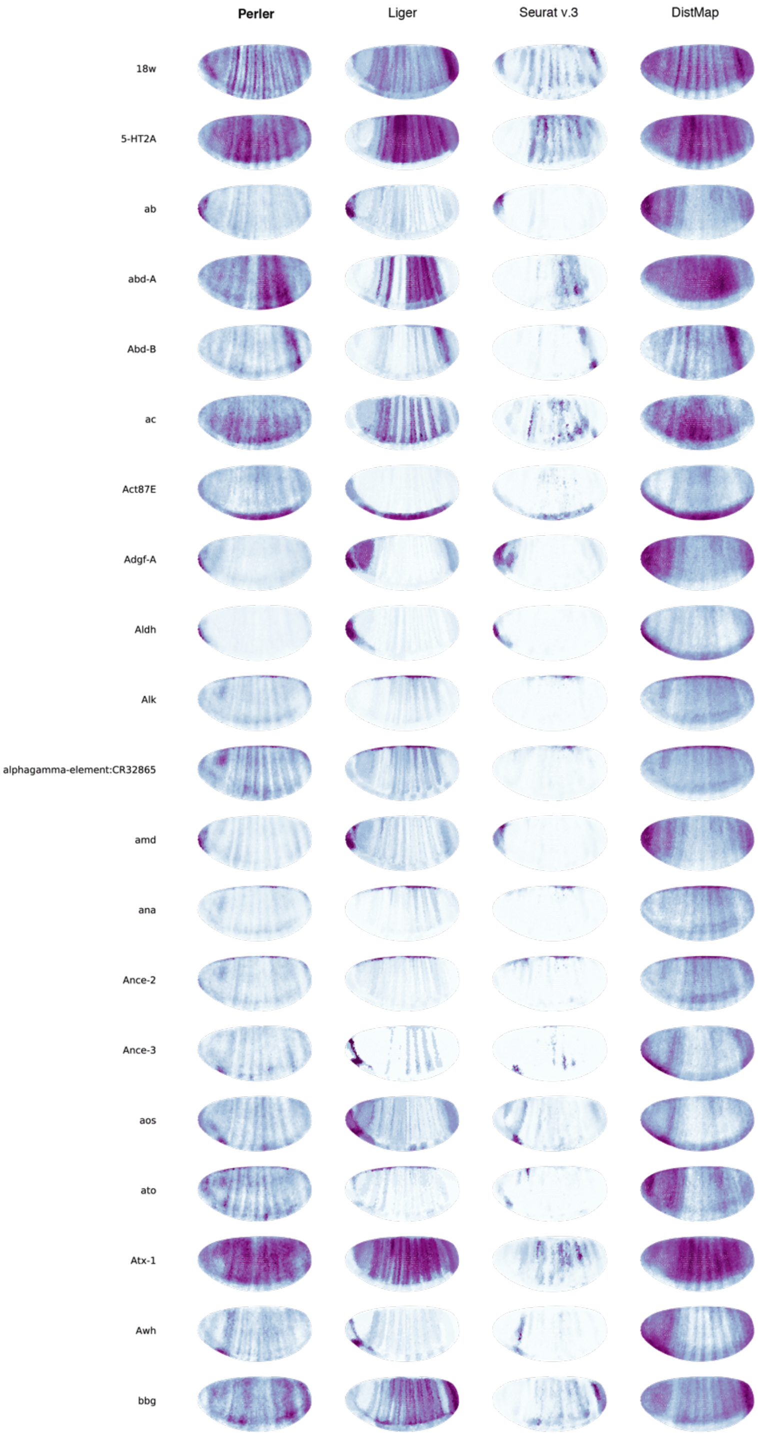


**Supplementary Figure 6: Spatial prediction of non-landmark spatially restricted genes (SRGs) (1)**

Predictions of expression of non-landmark SRGs (308 genes) were selected in Supplementary Fig. 5 by Perler, Liger, Seurat (v.3), and DistMap.

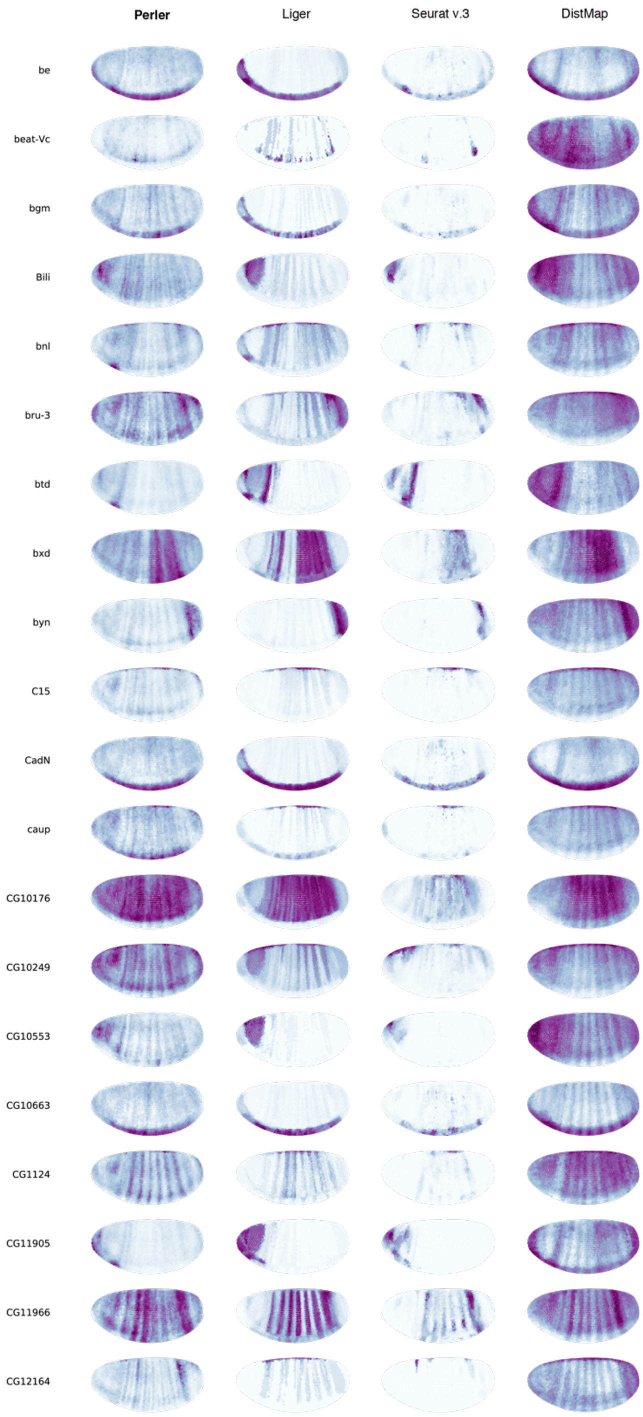


**Supplementary Figure 6 (2)**

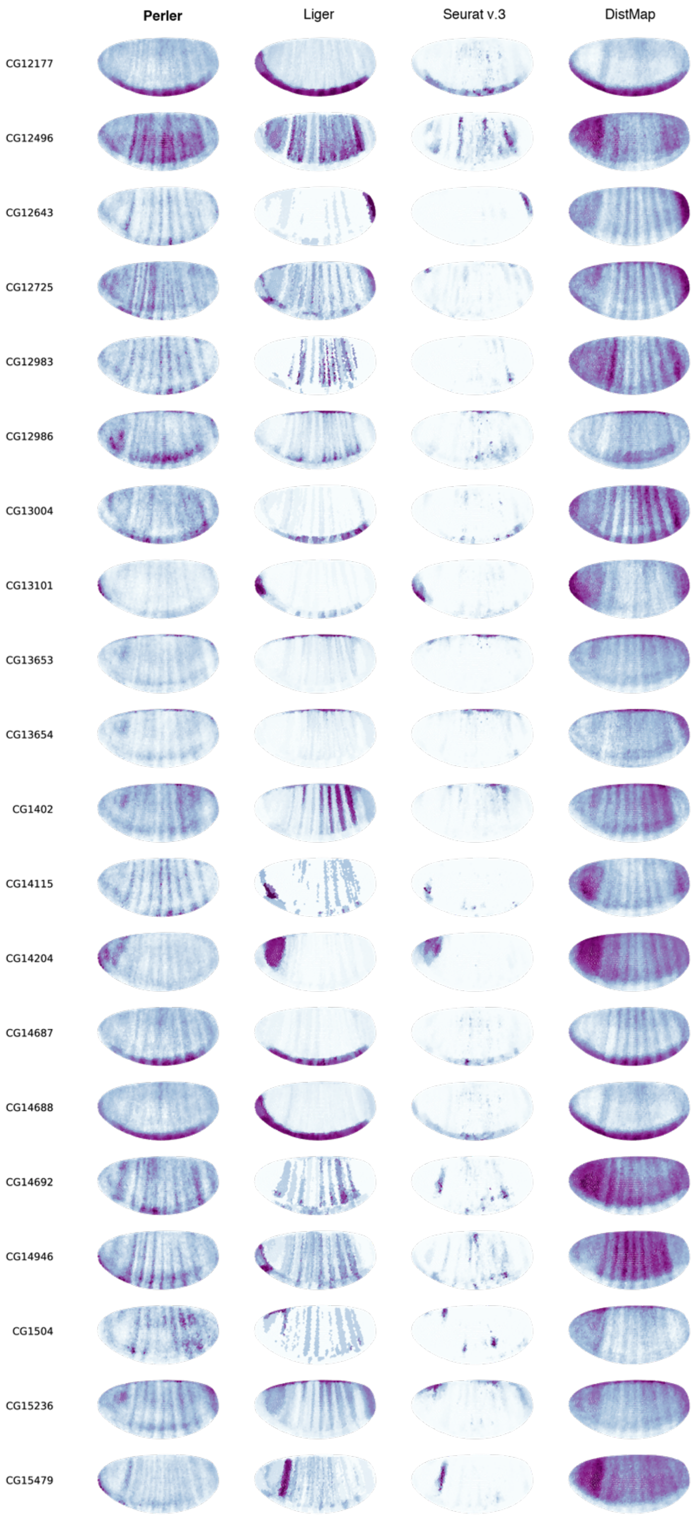


**Supplementary Figure 6 (3)**

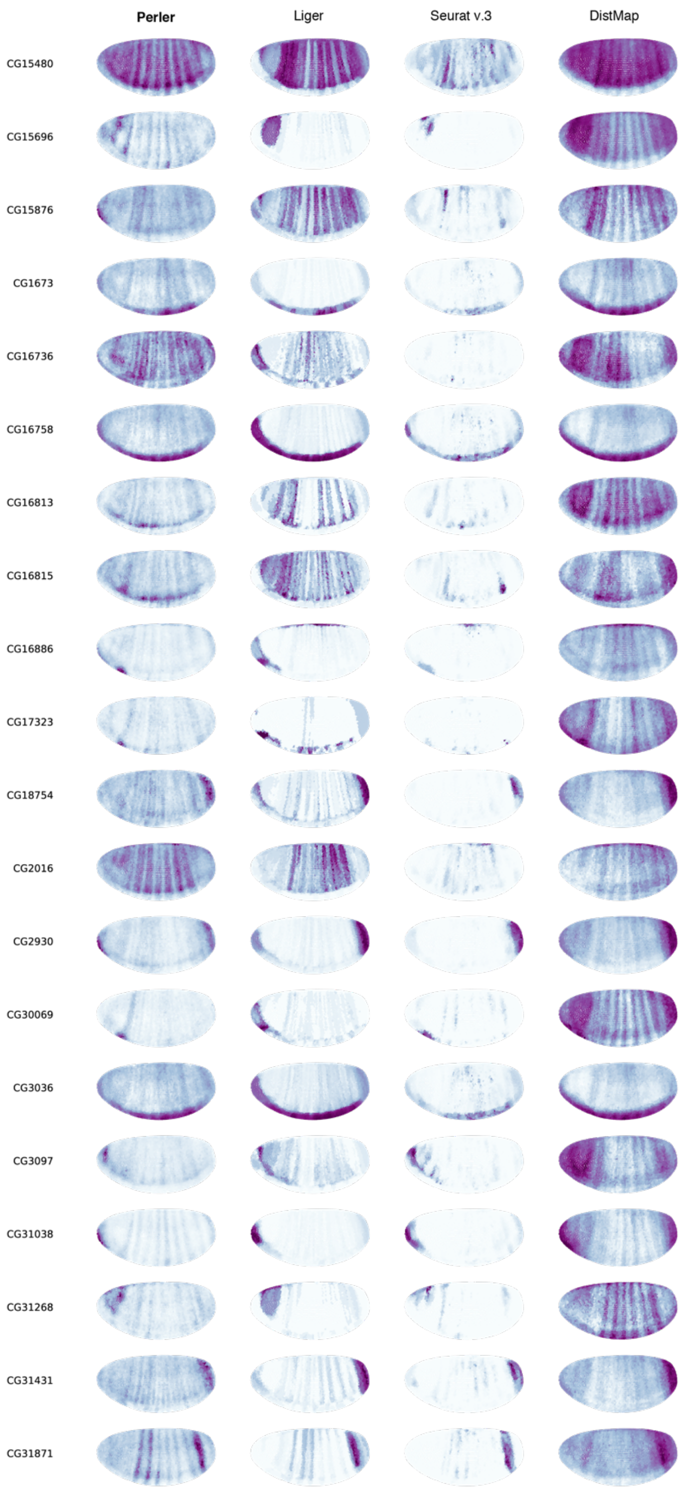


**Supplementary Figure 6 (4)**

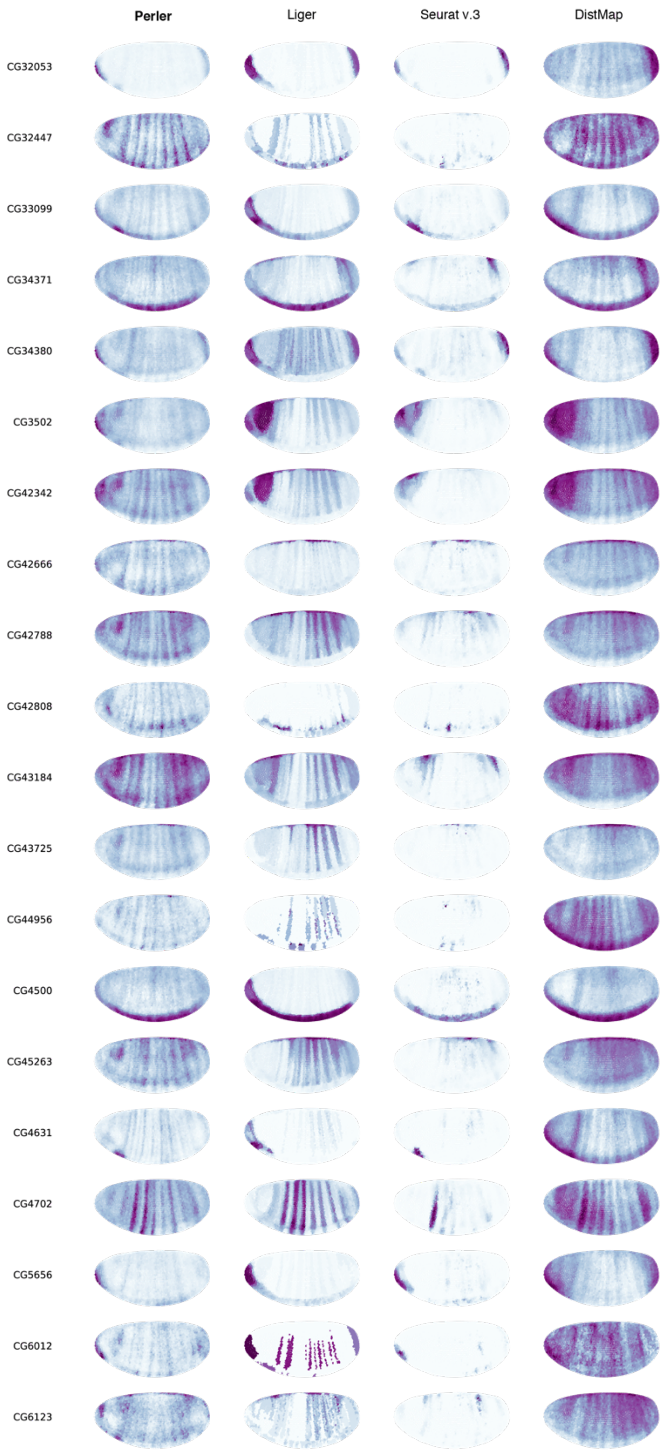


**Supplementary Figure 6 (5)**

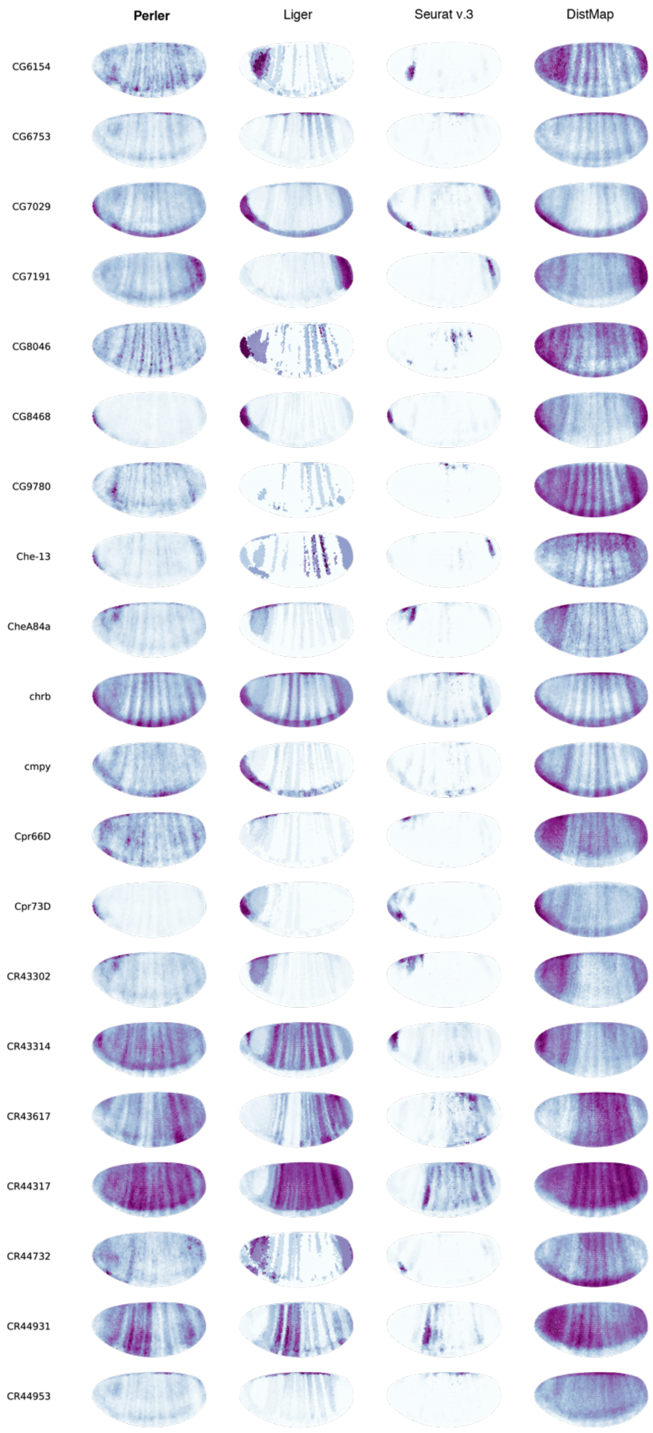


**Supplementary Figure 6 (6)**

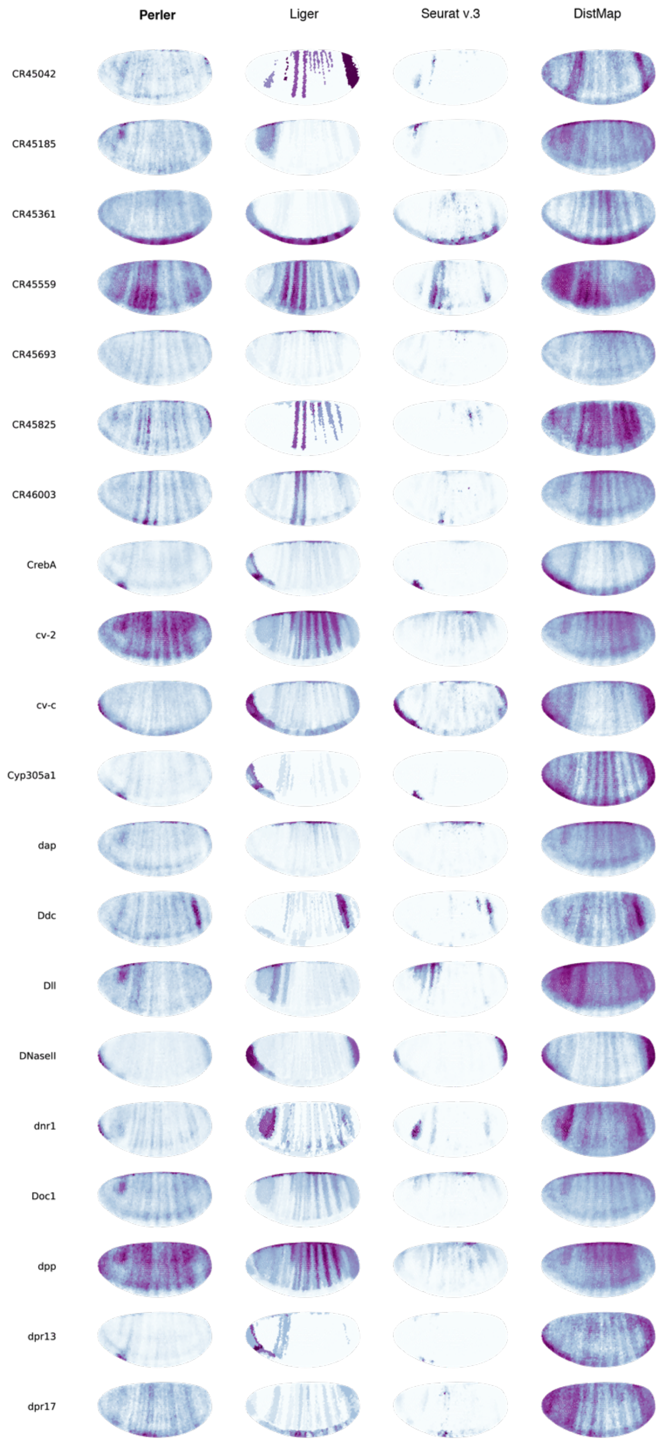


**Supplementary Figure 6 (7)**

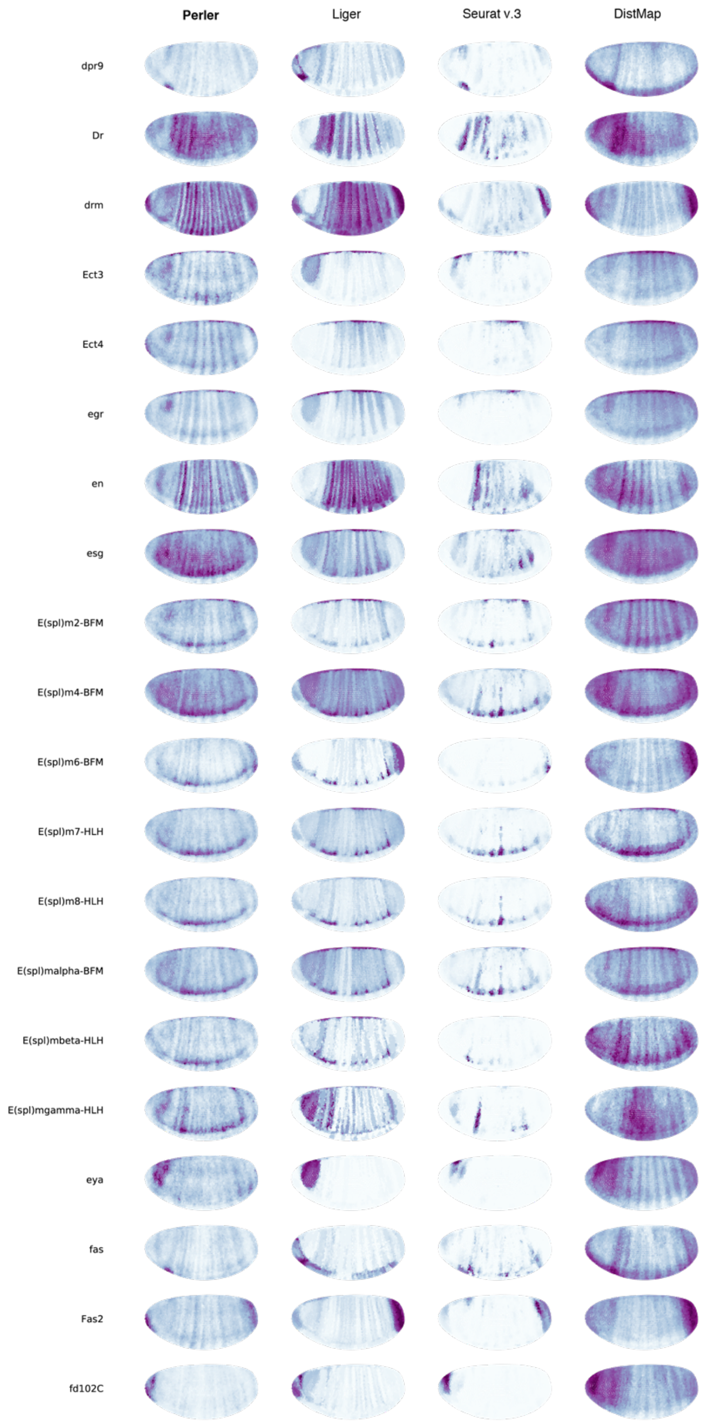


**Supplementary Figure 6 (8)**

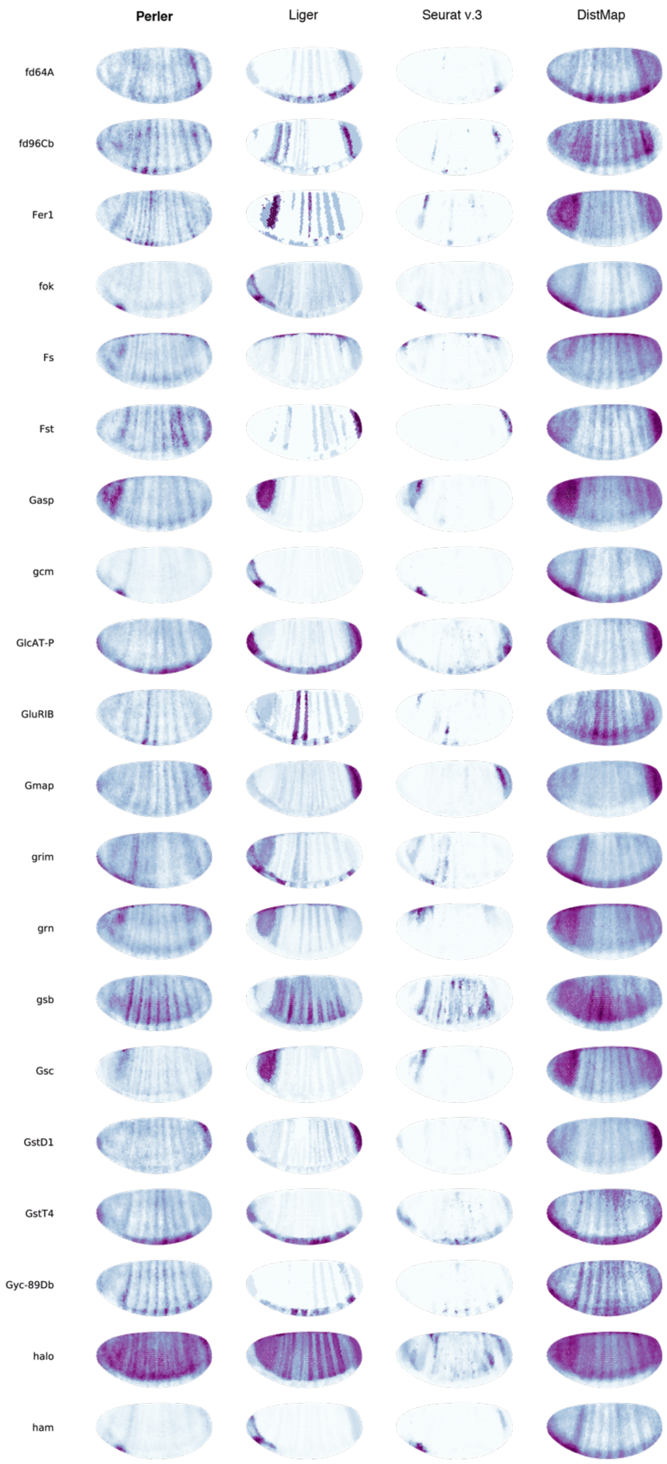


**Supplementary Figure 6 (9)**

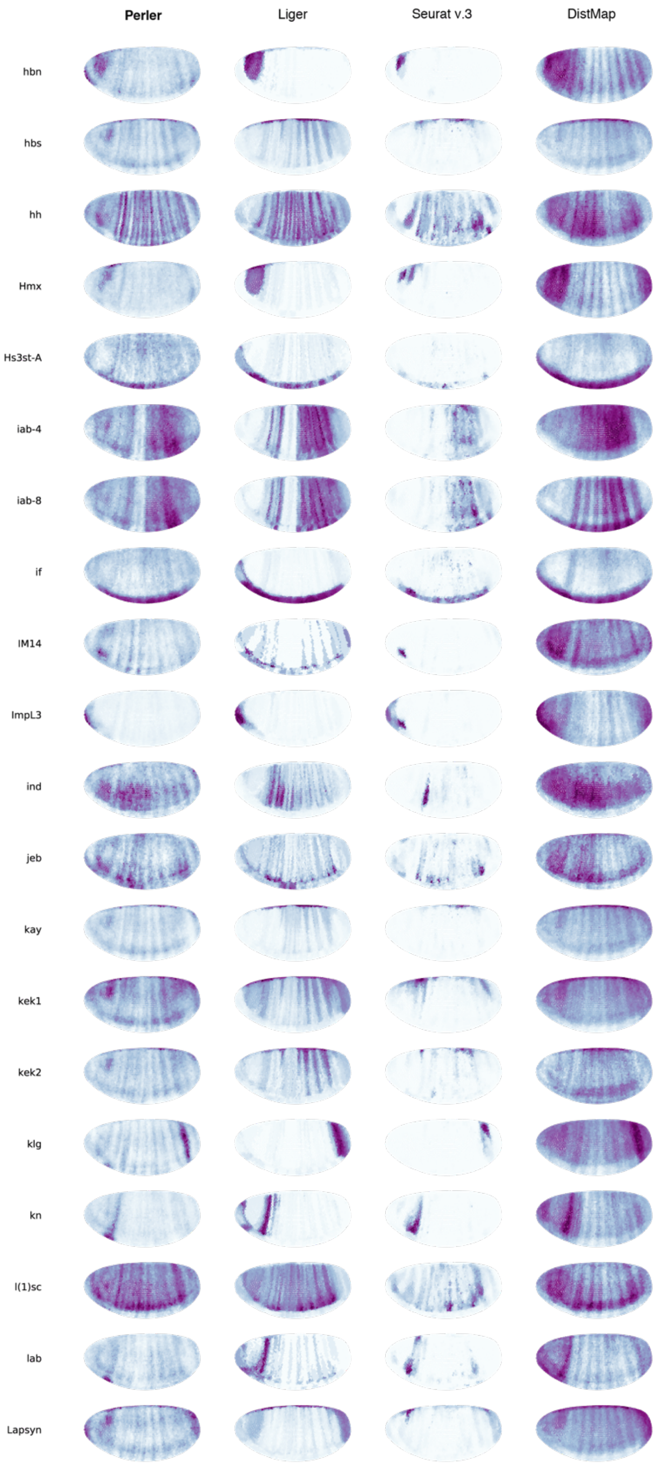


**Supplementary Figure 6 (10)**

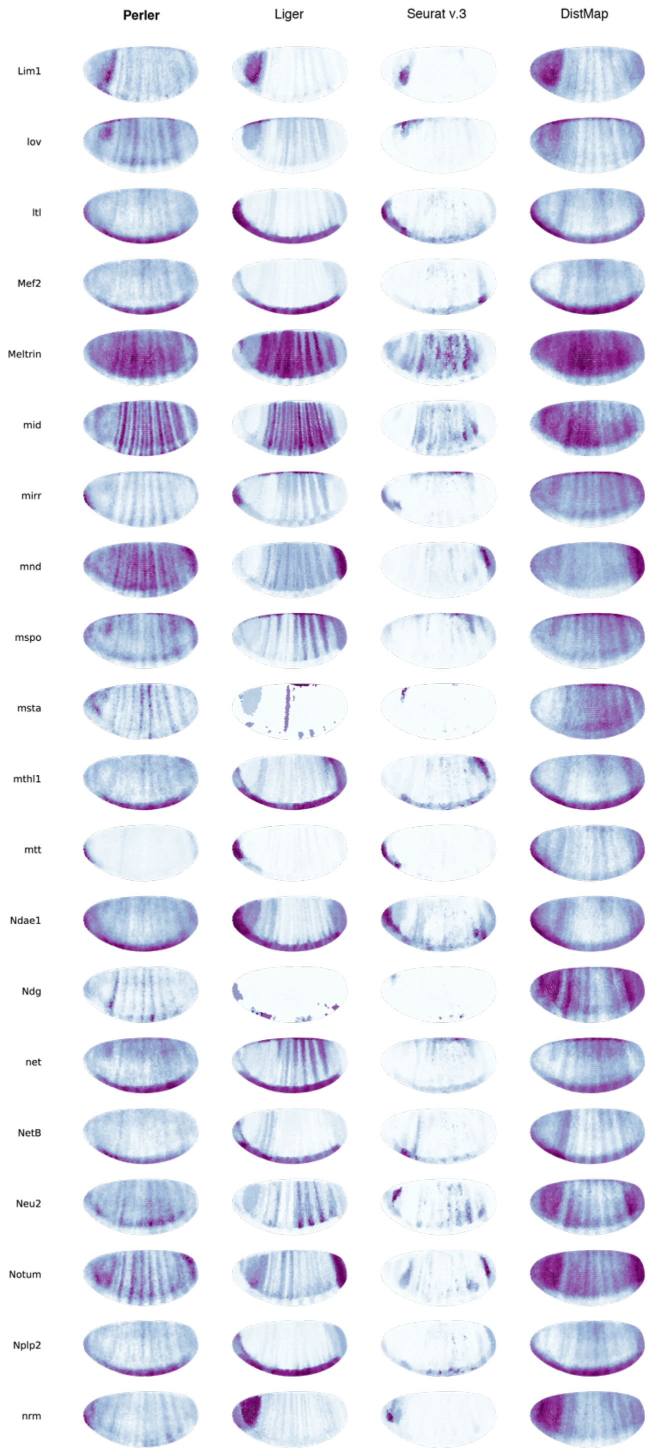


**Supplementary Figure 6 (11)**

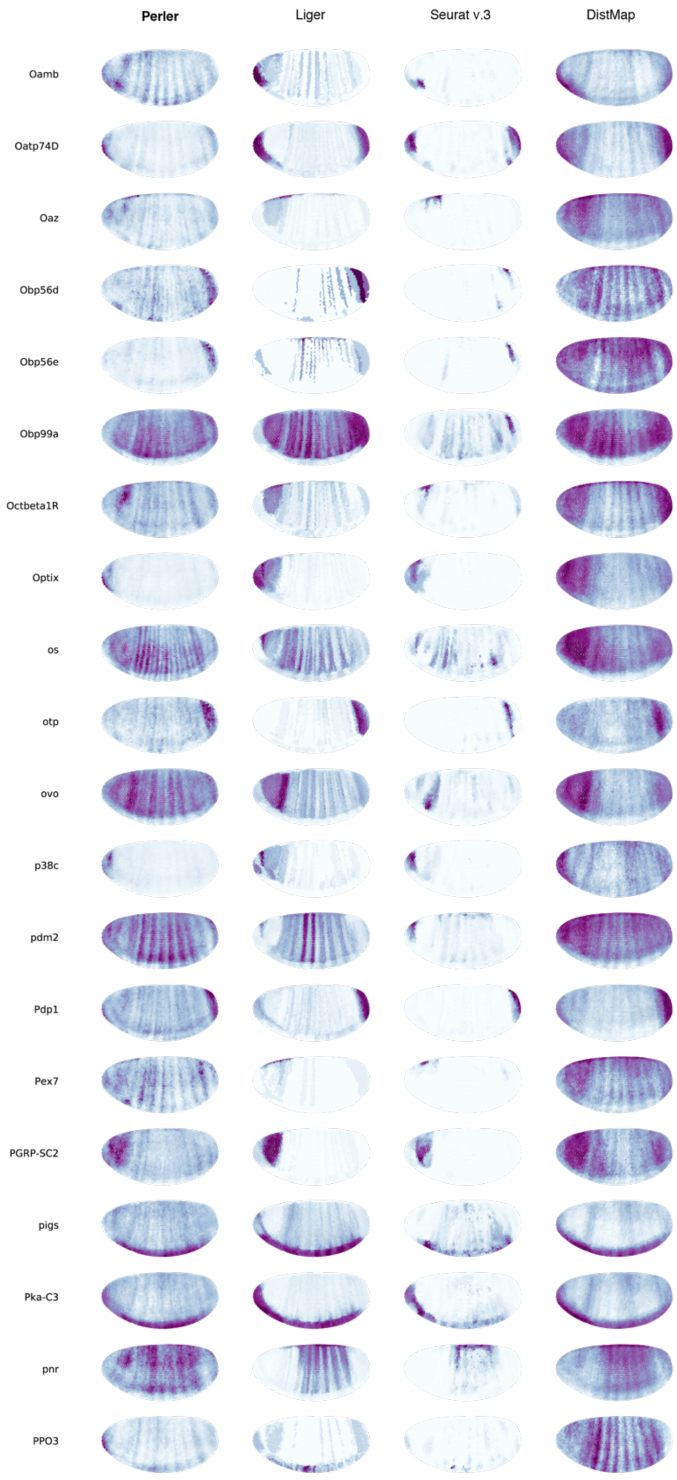


**Supplementary Figure 6 (12)**

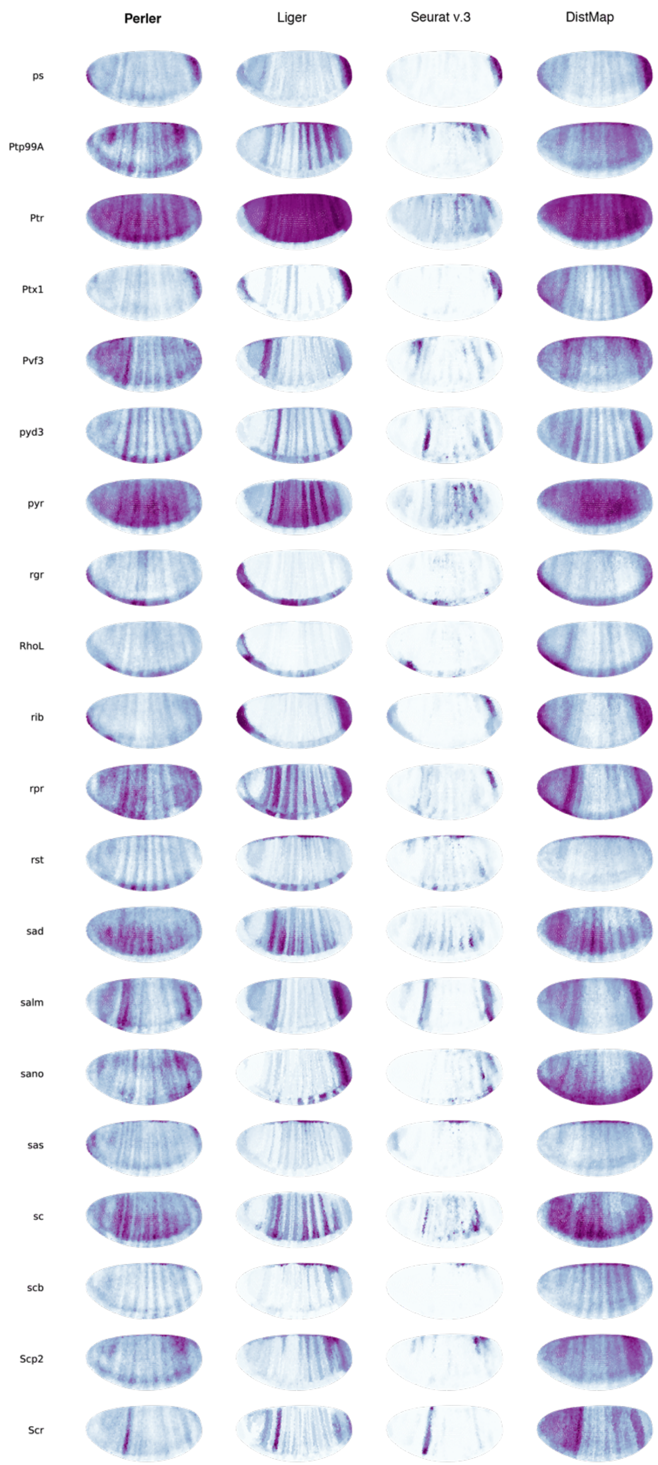


**Supplementary Figure 6 (13)**

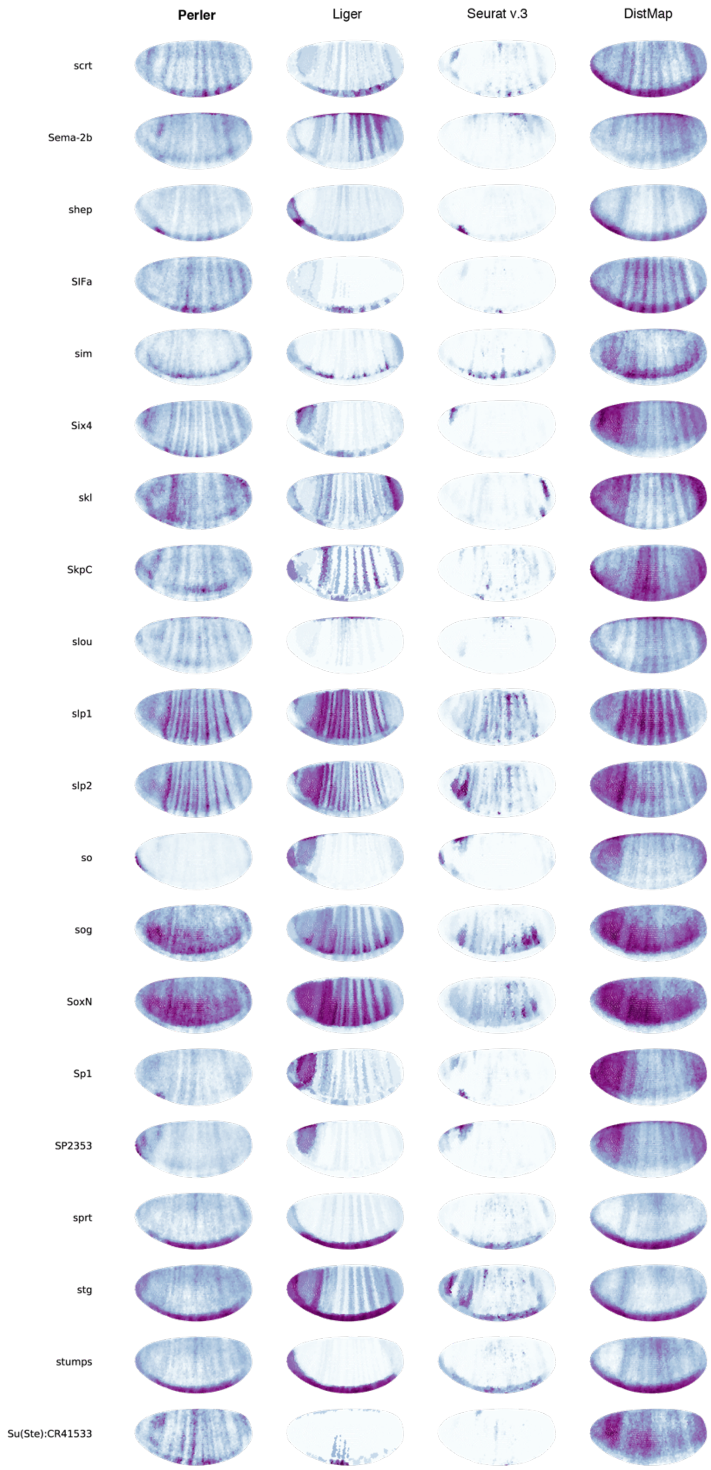


**Supplementary Figure 6 (14)**

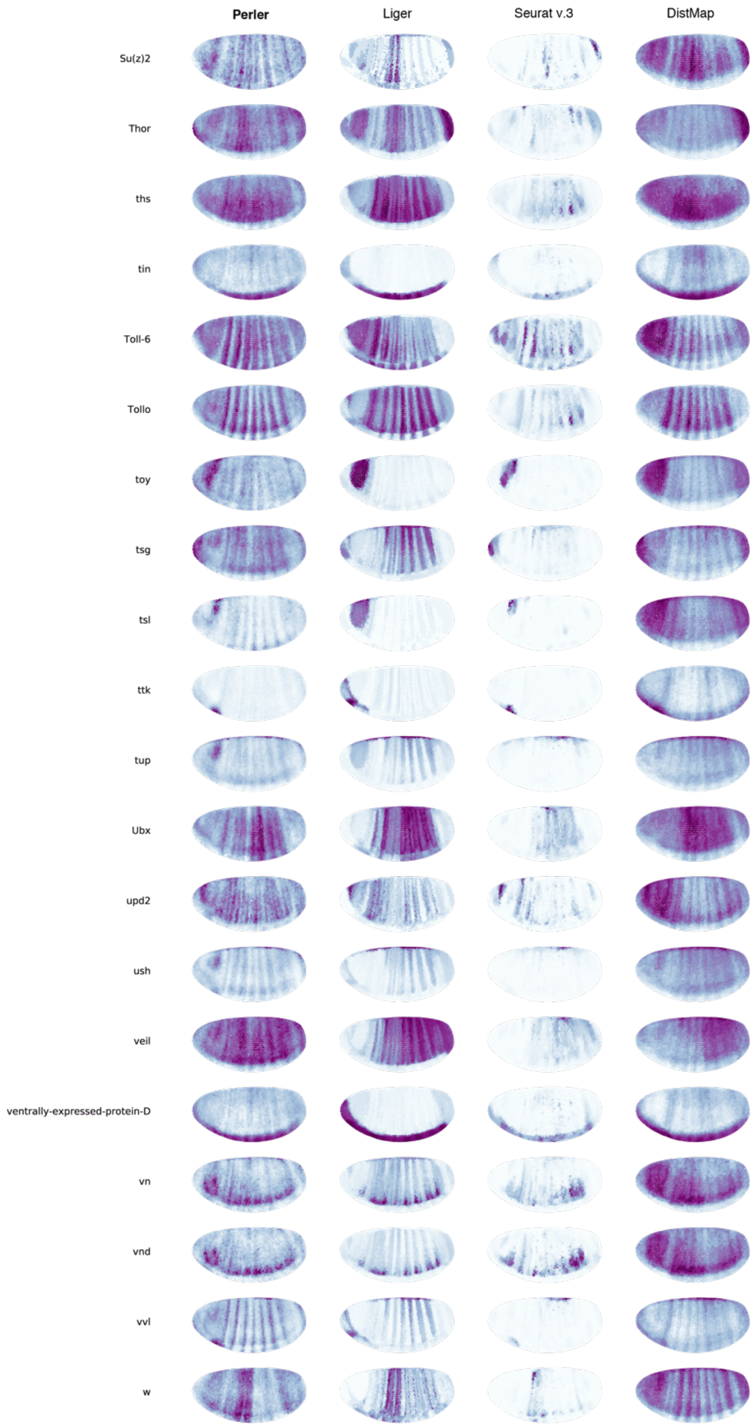


**Supplementary Figure 6 (15)**

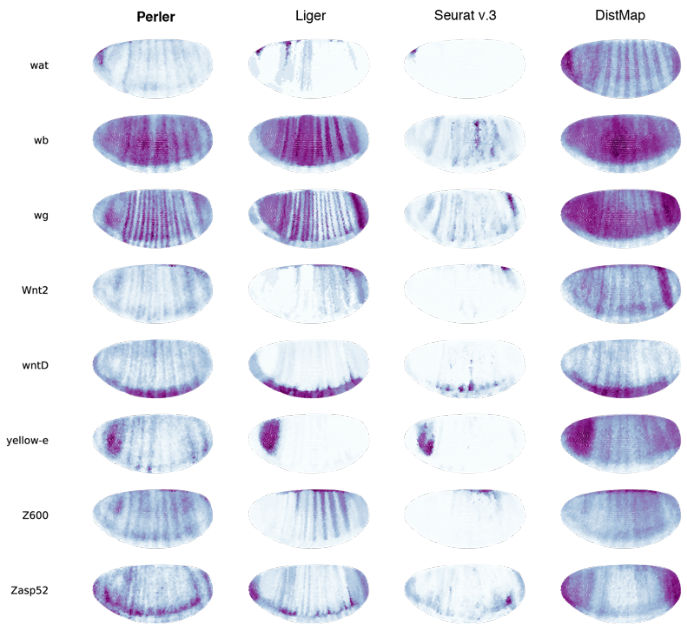


**Supplementary Figure 6 (16)**

**Supplementary Table 1: Comparison of Perler with existing methods**

| | Perler | Liger | Seurat v.3 | DistMap | Halpern et al,.[7] | Seurat v.1 |
|---|---|---|---|---|---|---|
| Continuous (not binary) | ✓ | ✓ | ✓ | × | ✓ | × |
| Applicability | ✓ | ✓ | ✓ | ✓[b] | × | ✓[b] |
| Dimensionality reduction | ✓[a] | ✓ | ✓ | × | × | × |
| Model-based mapping | ✓ | × | × | × | ✓ | ✓ |
| Generalization (not overfitting) | ✓ | × | × | × | - | - |

[a] Perler used a dimensionality reduction technique (PLSC) as a preprocessing

[b] DistMap and Seurat v.1 are applicable to the datasets whose ISH data is binarized

Perler characteristics relative to Liger, Seurat (v.3), DistMap, the method described by Halpern et al.[7], and Seurat (v.1).
